# Selective Vulnerability of Dopamine-Glutamate Neurons in Aging Weakens Entorhinal Dopamine Signaling

**DOI:** 10.1101/2025.10.06.680552

**Authors:** Jacquelyn N. Tomaio, Sixtine Fleury, Alexandra Bilder, Jordan Nacimba, Lakshman Abhilash, Yoon Seok Kim, Charu Ramakrishnan, Lief E Fenno, Karl Deisseroth, Susana Mingote

## Abstract

The lateral entorhinal cortex (LEC) supports novelty detection and episodic memory and is selectively vulnerable to aging. Dopamine signals novelty in LEC, but how aging alters this input is unclear. Using intersectional viral strategies to distinguish ventral tegmental area (VTA) dopamine neurons with or without glutamate co-release, we find that co-releasing neurons are a minority (∼30%) in the VTA, yet provide ∼93% of the dopaminergic projections to LEC. In aged mice, dopamine-glutamate labeling in VTA and LEC decreases ∼80%, whereas pan-dopaminergic labeling of LEC axons remains, consistent with functional silencing rather than degeneration. In LEC dopaminergic axons, dopamine synthesis is reduced while glutamate vesicular packaging is relatively spared. Optogenetic stimulation combined with a dopamine sensor reveals diminished dopamine release at high frequencies, when synthesis demand is greatest. These findings identify selective vulnerability of dopamine–glutamate neurons as a cell- and circuit-specific mechanism that weakens dopamine signaling within memory circuits.

## Introduction

Across species, aging is often accompanied by a gradual decline in cognitive abilities, with episodic memory among the most consistently affected ^1, 2^. Episodic memory supports the ability to remember specific events within their broader context, and when it weakens, everyday situations become harder to navigate ^3^. Older individuals may show a reduced response to novelty, greater difficulty distinguishing new from familiar events, and increased susceptibility to interference and false memories ^4–10^. These behavioral deficits are thought to reflect, in part, impairments in mnemonic discrimination, the ability to distinguish between similar experiences or memory representations, which depends on pattern separation processes ^2, 11^.

Age-related brain changes arise in the absence of overt neurodegeneration and are thought to reflect subtle, circuit-level dysfunction within medial temporal lobe networks that support memory ^12–15^. Within this network, the lateral entorhinal cortex (LEC) emerges as one of the earliest and most vulnerable regions in older adults with mild cognitive impairment and in preclinical Alzheimer’s disease ^16–20^. The LEC supports novelty detection, item-context integration, and associative memory ^21–26^, and reduced LEC engagement in older adults is linked to poorer novel object discrimination ^19^. However, the afferent inputs and neuromodulatory signals that contribute to the age-related attenuation of LEC responses to novel events remain unknown.

Dopamine neurons in the ventral tegmental area (VTA) signal the novelty and salience of incoming stimuli ^27–29^ and modulate novelty discrimination ^30, 31^. VTA dopaminergic innervation of the LEC is dense and patchy, forming dopaminergic islands ^32, 33^. Despite being far denser than dopaminergic input to the hippocampus ^34–37^, the functional role of this VTA-to-LEC pathway has received comparatively little attention. Only recently has work begun to define its contribution, providing evidence that dopamine in the LEC supports associative memory, in part by encoding the novelty status of cues ^38^. Aging is associated with reduced dopaminergic neuromodulation ^39^ that is not uniformly expressed across the forebrain. Human imaging studies indicate that declines in dopaminergic markers follow region-specific trajectories across the adult lifespan, suggesting selective vulnerability of particular dopamine circuits ^40–42^. Evidence for parallel dopamine circuits comes from rodent studies showing that VTA dopamine neurons comprise genetically and projection-defined subpopulations that differentially innervate functionally distinct regions ^43–48^, including medial temporal lobe memory regions ^33, 36, 47^. However, most work on age-related dopaminergic decline has centered on the striatum and prefrontal cortex, leaving open how aging reshapes dopaminergic input to early-vulnerable memory regions such as the LEC. This gap is clinically relevant because nonselective approaches that elevate dopamine signaling globally have produced variable cognitive effects ^49–52^, consistent with the idea that circuit-specific dopamine failures require circuit-specific interventions.

VTA neurons can be divided into distinct subpopulations based on their ability to co-release additional neurotransmitters ^53, 54^. Among these, dopamine-glutamate (DA-GLU) neurons represent a unique subpopulation ^55–59^, which exhibit strong responses to salient stimuli and contribute to behavioral flexibility ^30, 54, 56, 60–63^. DA-GLU neurons rely on the vesicular glutamate transporter 2 (VGLUT2) for glutamate loading and release ^58, 64^ and make strong glutamatergic connections to LEC neurons ^34^. Co-transmission from VTA dopamine neurons is increasingly recognized as sensitive to aging, with reduced tyrosine hydroxylase (TH) and VGLUT2 mRNA expression observed in aged mice ^65^. To determine how aging affects VTA dopamine neuron subpopulations and their dense projections to the LEC, we labeled pre-defined VTA dopamine subpopulations based on co-transmission profile and projection target using intersectional strategies ^66^. Through complementary anatomical and functional approaches, we identify selective vulnerability of dopamine–glutamate neurons as a cell type- and circuit-specific mechanism underlying the weakening of dopamine signaling within memory circuits.

## Results

### Labeling Subpopulations of Dopamine Neurons based on their Glutamate Co-Transmission Profile

To distinguish dopamine neurons that co-release glutamate (DA-GLU) from those that release only DA (DA-only), we used an INTRSECT 2.0 intersectional viral approach ^66^ in double-transgenic TH-Flp/+::VGLUT2-Cre/+ mice (**Fig. 1a, top**). We injected a mixture of two INTRSECT viruses unilaterally into the middle VTA and the lateral VTA/substantia nigra pars compacta to increase the chances of transfecting all VTA dopamine neurons (**Fig. 1a, middle**). AAV8-Con/Fon-EYFP labels neurons that co-express Flp and Cre (DA-GLU; EYFP+), whereas AAV8-Coff/Fon-mCherry labels Flp+ neurons lacking Cre (DA-only; mCherry+). In this design, “Con” denotes Cre-dependent activation of the fluorescent construct, whereas “Coff” denotes a Cre-off configuration in which Cre suppresses construct expression (**Fig. 1a, bottom**).

**Fig. 1.**
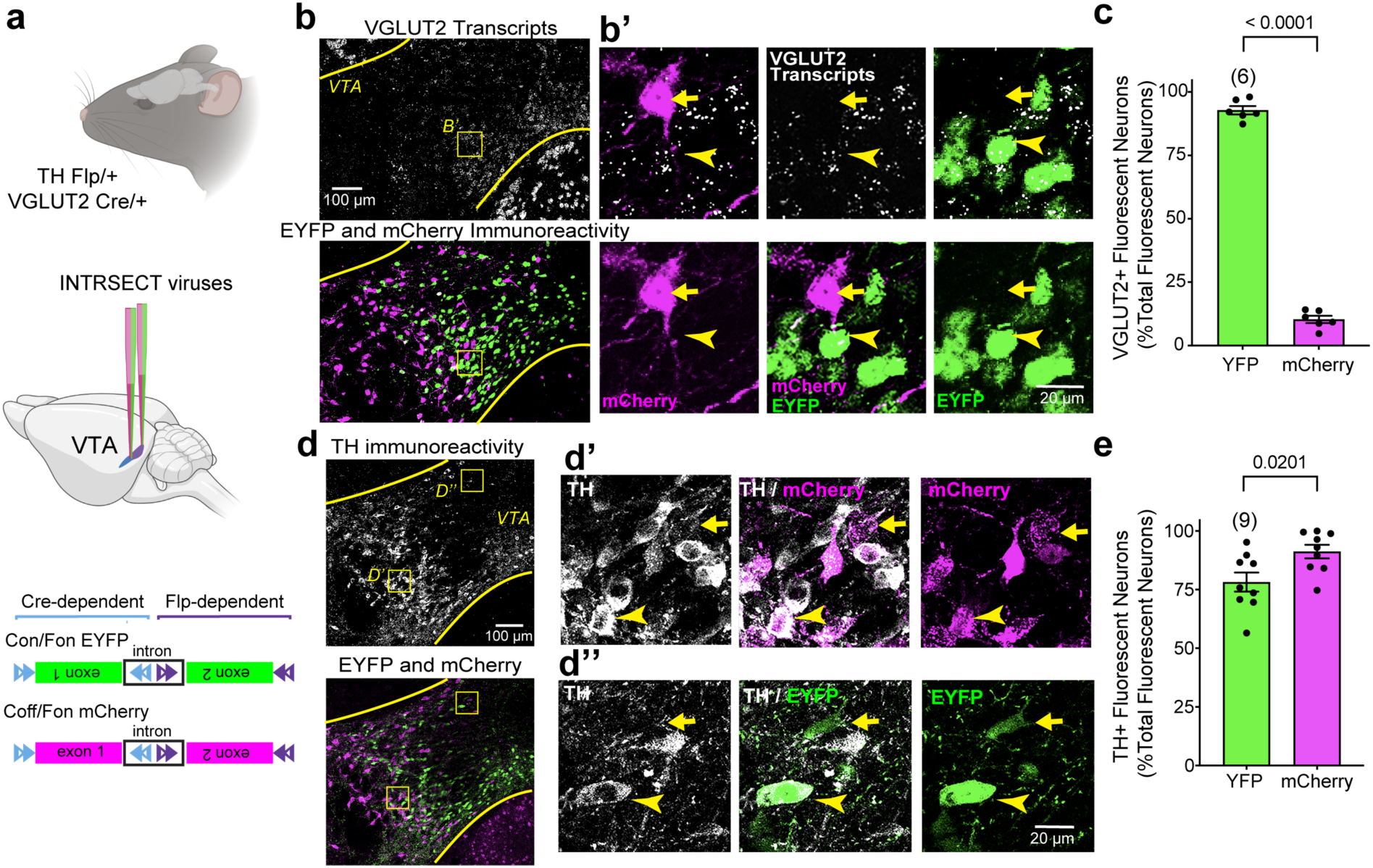
Intersectional labeling distinguishes DA-GLU and DA-only neurons in the VTA using INTRSECT viruses in young mice. **a**, Experimental schematic for double transgenic TH-Flp/+::VGLUT2-Cre/+ mice and INTRSECT constructs. Con/Fon drives EYFP in neurons that express both Flp and Cre (DA-GLU, shown in green) Coff/Fon drives mCherry in neurons that express Flp but not Cre (DA-only, shown in magenta); schematic adapted from Fenno et al., 2020. **b**, VTA images combining VGLUT2 (*Slc17a6*) mRNA *in situ* (top) with EYFP and mCherry immunostaining (bottom); Inset B′ shows EYFP+ somata containing VGLUT2 transcripts (arrowheads) and mCherry+ somata not (arrows). **c**, Quantification of fluorescent neuron subtypes co-expressing VGLUT2 mRNA and EYFP (green) or VGLUT2 mRNA and mCherry (magenta), expressed as the percentage of total fluorescently labeled neurons. EYFP+ neurons show higher VGLUT2 co-expression (Welch’s test, t=37.90, df=9.843). **d**, Confocal images of VTA sections showing fluorescent labeling of TH+ (top), and EYFP+ and mCherry+ neurons (bottom); Arrowheads in inset D′ indicate colocalization of TH and mCherry immunoreactivity; the arrow marks a rare cell that is mCherry+ but lacks TH immunoreactivity. Arrowheads in inset D″ indicate colocalization of TH and EYFP immunoreactivity; arrows mark the few EYFP+ neurons that do not express TH. **e,** Quantification of fluorescent neuron subtypes co-expressing TH protein and EYP (green) or TH protein and mCherry (magenta), expressed as the percentage of total fluorescently labeled neurons. The specificity is higher for mCherry expression (Welch’s test, t=2.612, df=14.48). Bar graphs show mean percentages ± SEM; number of mice shown in parentheses; *p* values shown above brackets.

We validated that INTRSECT expression required both recombinases (Flp and Cre) by injecting the same viral mixture into young TH-Flp/+ or VGLUT2-Cre/+ mice. Con/Fon-driven EYFP expression was absent in both single-transgenic lines, confirming that Con/Fon expression requires the presence of both Cre and Flp recombinases (**Extended Fig. 1a**). By contrast, Coff/Fon-driven mCherry expression was robust throughout the VTA in TH-Flp/+ mice, indicating that in the absence of Cre, the virus is activated by Flp alone and is therefore broadly expressed, including in midline VTA regions enriched in DA-GLU neurons. In VGLUT2-Cre/+ mice, however, mCherry was absent, consistent with suppression of the Coff/Fon construct by Cre (**Extended Fig. 1b**).

To assess glutamatergic identity and specificity, we combined VGLUT2 *in situ* hybridization with EYFP and mCherry immunohistochemistry. Confocal images (**Fig. 1b)** and volume renderings (**Extended Fig. 1c**) showed that 93% ± 1.6% of EYFP+ neurons contained VGLUT2 puncta, confirming successful targeting of DA–GLU neurons. mCherry+ neurons showed minimal VGLUT2 signal, although 10% ± 1.4% displayed a single VGLUT2 punctum (**Fig. 1c; Extended Fig. 1d**).

To evaluate dopaminergic specificity, we quantified co-localization with TH immunostaining. Overall, 91% ± 2.9% of mCherry+ neurons and 78% ± 4.1% of EYFP+ neurons were TH+ (**Fig. 1d,e**). The EYFP+ / TH− fraction may reflect a previously described subset of VTA neurons that express TH mRNA but have undetectable TH protein ^67^. Given that VGLUT2 signal is present in most EYFP+ neurons, these EYFP+ / TH− cells could represent glutamatergic-only neurons with ectopic EYFP expression.

Together, these data show that the INTRSECT strategy robustly labels DA-GLU neurons with EYFP and DA-only neurons with mCherry, enabling population-specific analyses of VTA circuits and their vulnerability to aging.

### Distribution of Subpopulations of Dopamine Neurons in the VTA and LEC in Young Mice

DA neuron subpopulations show distinct prevalence and spatial distribution patterns within the VTA (**Fig. 2a-c**). We found that 36% of all fluorescently labeled cells in the VTA were EYFP+, and when limiting the analysis to TH+ neurons, 33% of transfected TH+ VTA neurons were EYFP+ DA-GLU neurons (**Fig. 2c**). This proportion is consistent with prior reports in mice, which estimate DA-GLU neurons comprise 20–30% of the VTA DA population using intersectional genetic labeling or multiplex *in-situ* hybridization ^47, 54, 56, 67, 68^. We observed no differences between males (n=5) and females (n=4); therefore, the data were combined (**Extended Fig. 2a-c**).

**Fig. 2.**
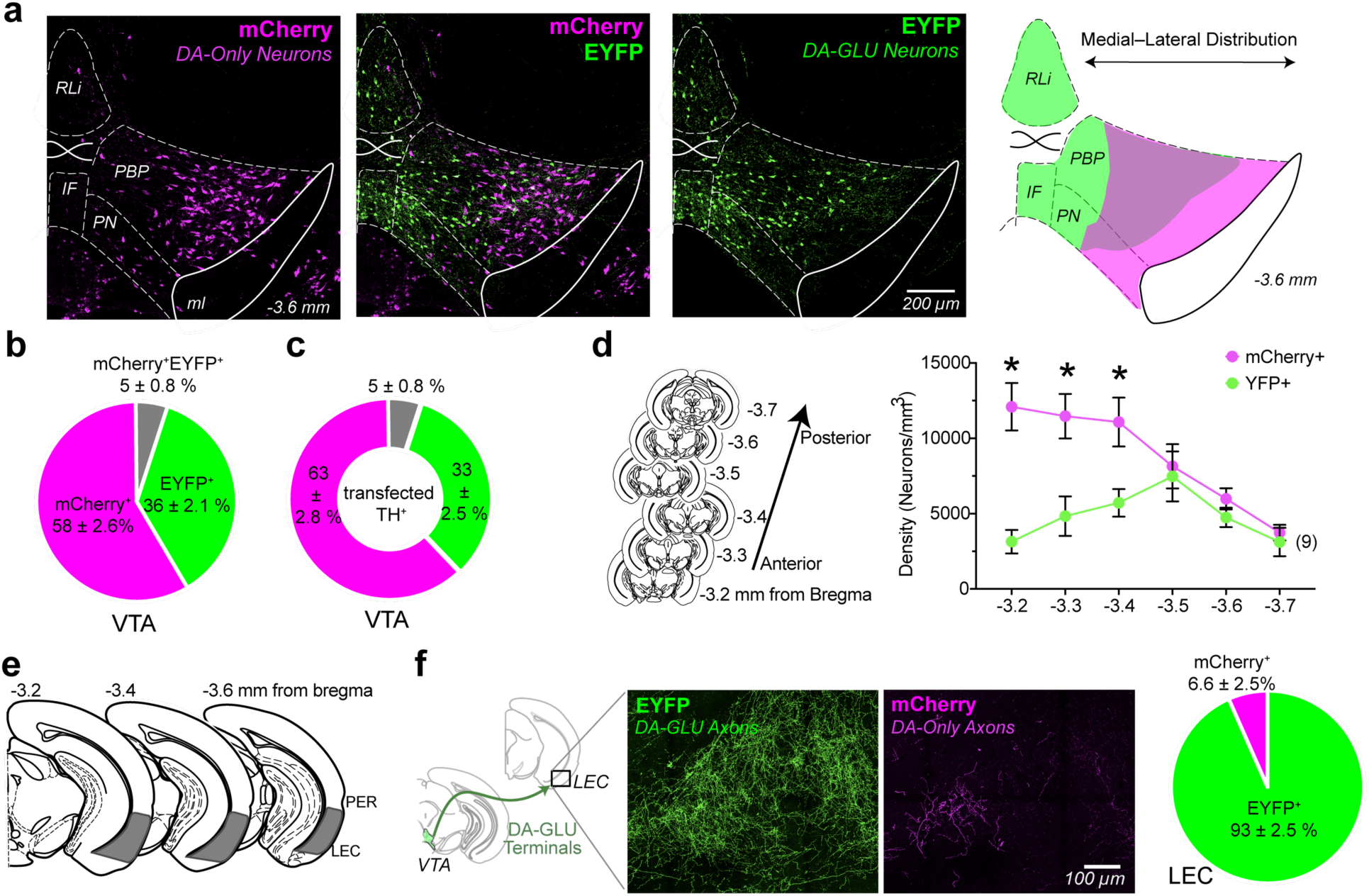
DA-GLU and DA-only neurons exhibit distinct spatial organization in the VTA and differentially innervate the LEC in young mice. **a**, Confocal photomicrographs showing the distribution of INTRSECT labeled VTA neurons expressing EYFP (DA-GLU, green), mCherry (DA-only, magenta), and merge (middle); schematic of the medial to lateral subtype distribution on the right. **b**, Proportion of EYFP+ (green), mCherry+ (magenta), and co-localized EYFP+/ mCherry+ (gray) neurons in the VTA as a percentage of all INTRSECT-labeled neurons. **c**, Proportion of EYFP+/TH+ (green), mCherry+/TH+ (magenta), and EYFP+/ mCherry+/TH+ (gray) neurons in the VTA as a percentage of all TH+ INTRSECT-labeled neurons. **d**, Schematic of coronal brain sections along the anterior–posterior VTA axis (−3.2 to −3.7 mm from bregma) used for quantification (left); Line plot showing the density (neurons/mm³) of mCherry+ and EYFP+ neurons across the anterior–posterior VTA axis in mm from bregma (two-way repeated measures ANOVA showed a significant interaction: *F*(5, 40) = 5.58, *p* < .001; large effect size: ηp² = .24; *significant Post-hoc Bonferroni comparisons, p < 0.05). **e**, Schematic of coronal brain sections from −3.2 mm to −3.6 mm relative to bregma, highlighting the LEC (gray) below the perirhinal cortex (PER). **f**, Schematic of VTA DA-GLU projections to the LEC (left). Representative confocal images of DA-GLU axons (EYFP+, green) and DA-only axons (mCherry+, magenta) in the LEC (middle). Pie chart shows the proportion of fluorescent axons in the LEC that are EYFP+ (green) versus mCherry+ (magenta) (right). All data shows mean density ± SEM; number of mice shown in parentheses. *IF*, interfascicular nucleus; *ml*, medial lemniscus; *PBP*, parabrachial pigmented nucleus*; PN*, paranigral nucleus; *RLi*, rostral linear nucleus.

DA-GLU and DA-only neurons also differ in their anatomical distribution. EYFP+ DA-GLU neurons are enriched in the interfascicular nucleus (IF), rostral linear nucleus (RLi), medial paranigral nucleus (PN), and medial parabrachial pigmented nucleus (PBP) of the VTA ^68^. Along the anterior-posterior axis (−3.2 to −3.7 mm from bregma), DA-GLU neurons were predominantly located in the middle portion of this range, whereas DA-only neurons were more concentrated in the anterior VTA (**Fig. 2d**). Statistical analyses confirmed this difference; mCherry+ neuron density was significantly greater than EYFP+ density at −3.2, −3.3, and −3.4 mm from bregma (**Fig. 2d**).

When examining dopaminergic projections to LEC, a region known to be especially vulnerable to aging ^11, 17, 19, 69^, we found that 93% of labeled axons were EYFP+ (**Fig. 2e–f**), indicating that nearly all DAergic input from the VTA to the LEC originates from DA-GLU neurons. This was a surprising and novel finding, given the DA-GLU neurons constitute only a minority population within the VTA. However, it is consistent with our earlier functional connectome study showing robust glutamatergic input from DA neurons to LEC neurons ^34^. Together, these results validate that DA-GLU neurons likely play a dominant role in modulating LEC function.

In summary, intersectional strategies reveal that DA-GLU neurons make up roughly one-third of VTA DA neurons, are concentrated in the medial portion of VTA, and provide the majority of DAergic innervation to the LEC.

### Effects of Aging on the Density of Dopamine Neuron Subpopulations within the VTA

Aging affects dopaminergic markers heterogeneously, suggesting that distinct dopamine circuits show differential vulnerability ^40, 42, 70, 71^. Because DA-only and DA-GLU subpopulations innervate distinct brain regions ^54^, we investigated whether aging differentially impacts these specific subtypes.

We used the same INTRSECT viral strategy described above to label DA-GLU and DA-only neurons in mice at different ages: 3 months (young, n=9), 14 months (middle, n=7), and 24 months (aged, n=5) (**Fig. 3a**). We injected double mutant mice with INTRSECT viruses one month before reaching the target age to allow sufficient viral expression before perfusion. Age did not affect the viral transduction efficiency (∼75%), quantified as the percentage of TH+ neurons in the VTA that expressed either transgenes (mCherry and YFP combined) (**Fig. 3b**). Aging also did not alter the specificity of INTRSECT viruses in targeting DA-GLU versus DA-only neurons (**Extended Fig. 3a,b**). Having established that INTRSECT can be used to assess age-related differences, we then measured neuronal density of the labeled subpopulations. DA-only neurons showed a decreasing trend that did not reach statistical significance, whereas the density of DA-GLU neurons was significantly reduced by 14 months and declined further by 24 months, reaching an overall ∼60% decrease (**Fig. 3c**).

**Fig. 3.**
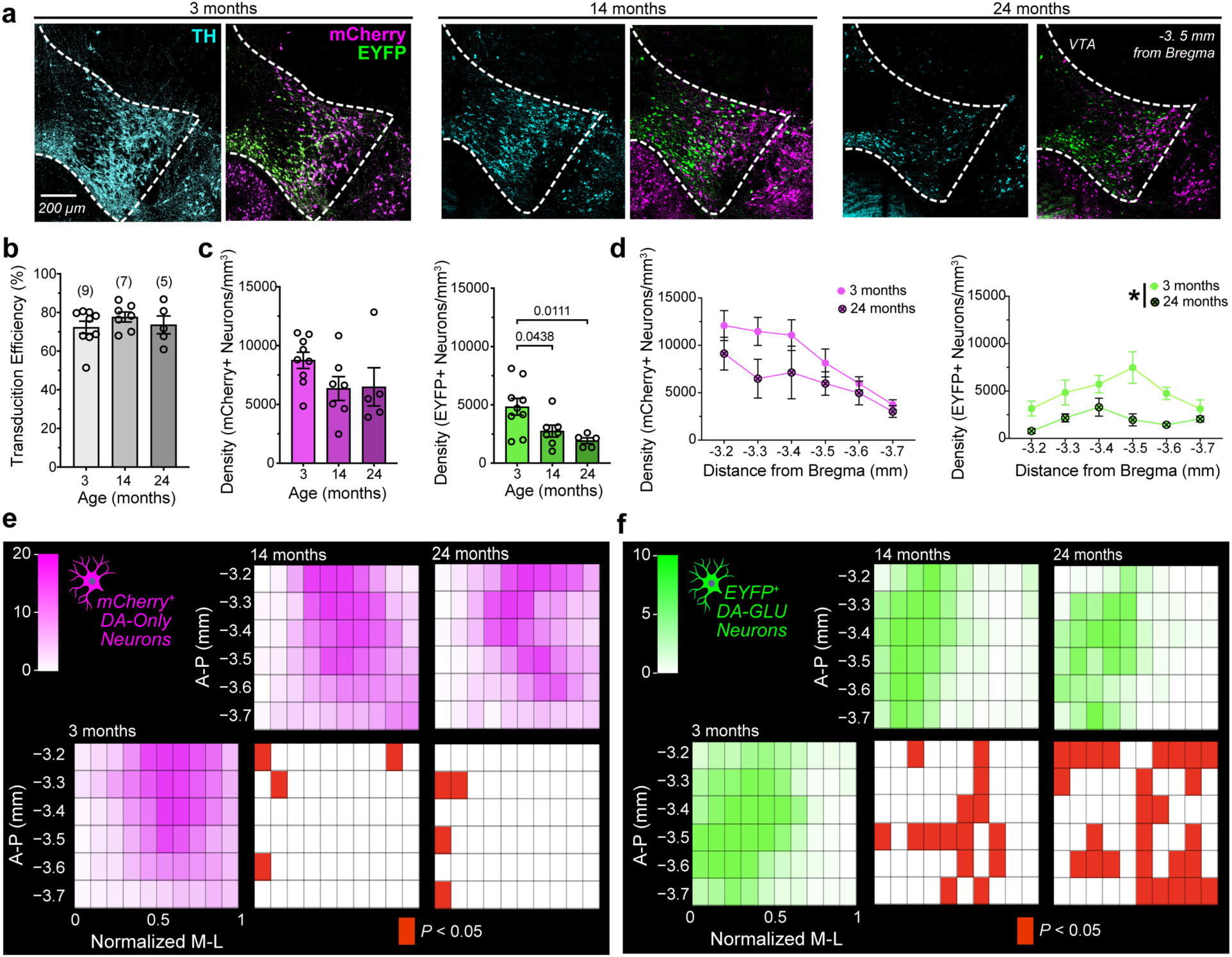
The effects of aging on the density and spatial distribution of dopamine neuron subpopulations within the VTA. **a**, Representative unilateral coronal sections through the VTA at approximately −3.5 mm from bregma in 3- (left), 14- (middle), and 24-month-old (right) mice, showing fluorescent labeling of TH+ (blue), EYFP+ (green), and mCherry+ (magenta) neurons. **b**, Bar graph showing the transduction efficiency, calculated as the percentage of TH+ neurons that co-expressed both EYFP and mCherry immunofluorescence, relative to the total number of TH+ VTA neurons, across age groups (no main effect of age: one-way ANOVA, F_(2, 18)_ = 8.271). **c**, Bar graph showing mCherry+ (left) and EYFP+ (right) neuronal density across age groups. Density of mCherry+ neurons shows no one-way ANOVA main effect of age (*F*_(2, 18)_ = 1.95, *p* = 0.171; moderate effect size: ηp² = 0.17). Density of EYFP+ neurons showed a main effect of age (F(2,18) = 5.81, p = 0.011; large effect size, ηp² = 0.39). Dunnett’s post hoc comparisons versus 3 months revealed significant differences in both the 14- and 24-month-old groups (exact *P* values are shown above the brackets). **d**, Line graphs showing the density of mCherry+ (top) and EYFP+ (bottom) neurons across six anterior–posterior VTA distances (−3.2 to −3.7 mm from bregma) in 3- and 24-month-old mice. Two-way repeated-measures ANOVA of mCherry+ neuronal density revealed a main effect of distance from bregma (F(5,60) = 8.04, p < 0.001, ηp² = 0.455), with no main effect of age and no age × distance interaction. For EYFP+ neuronal density, there was a main effect of age, indicated by an asterisk in the figure legend (F(1,12) = 8.29, p = 0.0138, ηp² = 0.11), as well as a main effect of distance from bregma (F(5,60) = 2.81, p = 0.024, ηp² = 0.25), with no significant interaction. **e,** 2D VTA density map showing the distribution of mCherry+ (DA-only) cells across anterior–posterior (A–P) and normalized medial–lateral (M–L) coordinates. Red tiles indicate bins that differed from the 3-month group (one-tailed *t*-test against the mean of the 3-month group), with p-values Benjamini–Hochberg corrected to control the false discovery rate (FDR) at 5%; only adjusted P < 0.05 bins are shown. **f**, Same as e, for EYFP+ (DA-GLU) cells. For other plots, data are mean ± SEM; numbers in parentheses indicate the number of mice.

We evaluated how density changes along the anterior–posterior axis by comparing 3- and 24-month-old mice (**Fig. 3d**). Density did not vary significantly along the anterior-posterior axis for the mCherry+ DA-only subpopulation. In contrast, aged (24-month-old) mice showed lower EYFP+ DA-GLU neuron density than young (3-month-old) mice across the anterior–posterior sections sampled. However, this analysis does not account for the significant heterogeneity in distribution of the subpopulations across the medial-lateral VTA axis (**Fig. 2a**). To further characterize age-related changes in cell-type distribution, we generated spatial density heat maps across the anterior-posterior and medial-lateral axes (**Fig.3e,f**). The maximum distance of TH+ cells from the midline was used as an estimate of the VTA medial-lateral dimension. This distance was greater in young males than in young females, and greater in middle-aged compared to young mice (**Extended Fig. 3c,d**), indicating that the medial-lateral VTA size varies with sex and age. To enable comparisons of cell distributions across age groups, we normalized each cell’s distance from the midline to the maximum distance measured in that section (i.e., expressed as a fraction of the longest distance). Comparing distributions across age groups revealed that DA-GLU neurons underwent the most pronounced spatial reorganization with aging, whereas DA-only neurons changed only modestly (**Fig. 3e,f**). In aged mice, reductions in DA-GLU cell counts were not uniform, with the greatest losses observed at the medial anterior (−3.2 mm) and posterior (−3.6 mm) levels, as well as more widespread losses across lateral regions. As a result, the remaining DA-GLU neurons in aged mice became concentrated within a medial VTA “hot spot” spanning approximately −3.3 to −3.5 mm relative to bregma. Consistent with this pattern, a DA-GLU minus DA-only difference map showed clear spatial segregation in young mice (ventromedial DA-GLU enrichment and lateral DA-only enrichment) that progressively collapsed with age, yielding widespread DA-only predominance and supporting preferential vulnerability of DA–GLU neurons and region-specific loss of their spatial niches (**Extended Fig. 3e)**.

In addition to a change in INTRSECT-labeled subpopulations, we also observed a decrease in the density of TH+ neurons in both middle-aged and aged mice (**Extended Fig. 3f**). The spatial mapping revealed little regional redistribution, with few bins showing significant age differences, consistent with a broadly uniform reduction rather than focal loss. Reduced TH expression could contribute to the loss of EYFP+ labeling, because INTRSECT expression depends on TH promoter activity in addition to VGLUT2 promoter activity. However, the spatial pattern of EYFP+ changes did not mirror the TH+ map, showing that the reduction in labeling cannot be explained solely by decreased TH expression.

Overall, our data identify DA-GLU neurons as a selectively vulnerable VTA subpopulation, with INTRSECT-labeling reduced by 14 months and further diminished in aged (24 months) mice. Spatial mapping indicates that this loss is clustered rather than uniform, suggesting that specific dopaminergic pathways may be preferentially affected, given the medial–lateral topography of VTA forebrain projections ^45^. Finally, the age-related spatial changes in EYFP+ DA-GLU neurons did not mirror those of TH+ cells, indicating that the reduction in labeling cannot be explained solely by decreased TH expression and raising the possibility that diminished VGLUT2 expression, together with reduced TH, may produce a “double hit” that contributes to the loss of DA-GLU labeling.

### Effects of Aging on the Density of Dopaminergic Projections to LEC

Since DA-GLU neurons provide the predominant dopaminergic input to the LEC, we next asked whether their age-related decline was accompanied by a corresponding reduction in INTRSECT-labeled axons within this region (**Fig. 4a**). To quantify terminal density, we performed 3D reconstructions of INTRSECT-labeled axons using a volume-rendering software and measured their volume relative to the total volume of the LEC. We observed a marked reduction (∼75%) in the density of EYFP+ DA-GLU axons in the LEC in both middle-aged and aged mice (**Fig. 4b**). In some aged animals, EYFP labeling was nearly absent, as illustrated in the representative photomicrograph (top right, **Fig. 4a**). We also examined the smaller population of mCherry+ DA-only axons in the LEC and found that their density was also significantly reduced at 14 and 24 months (**Fig. 4c**). Although DA-GLU neuron density in the VTA declined by only ∼40% at 14 months, INTRSECT-labeled axons in the LEC were reduced by nearly 75%. This disproportionate decline suggests that LEC dopaminergic input from DA–GLU neurons is especially vulnerable early in aging.

**Fig. 4.**
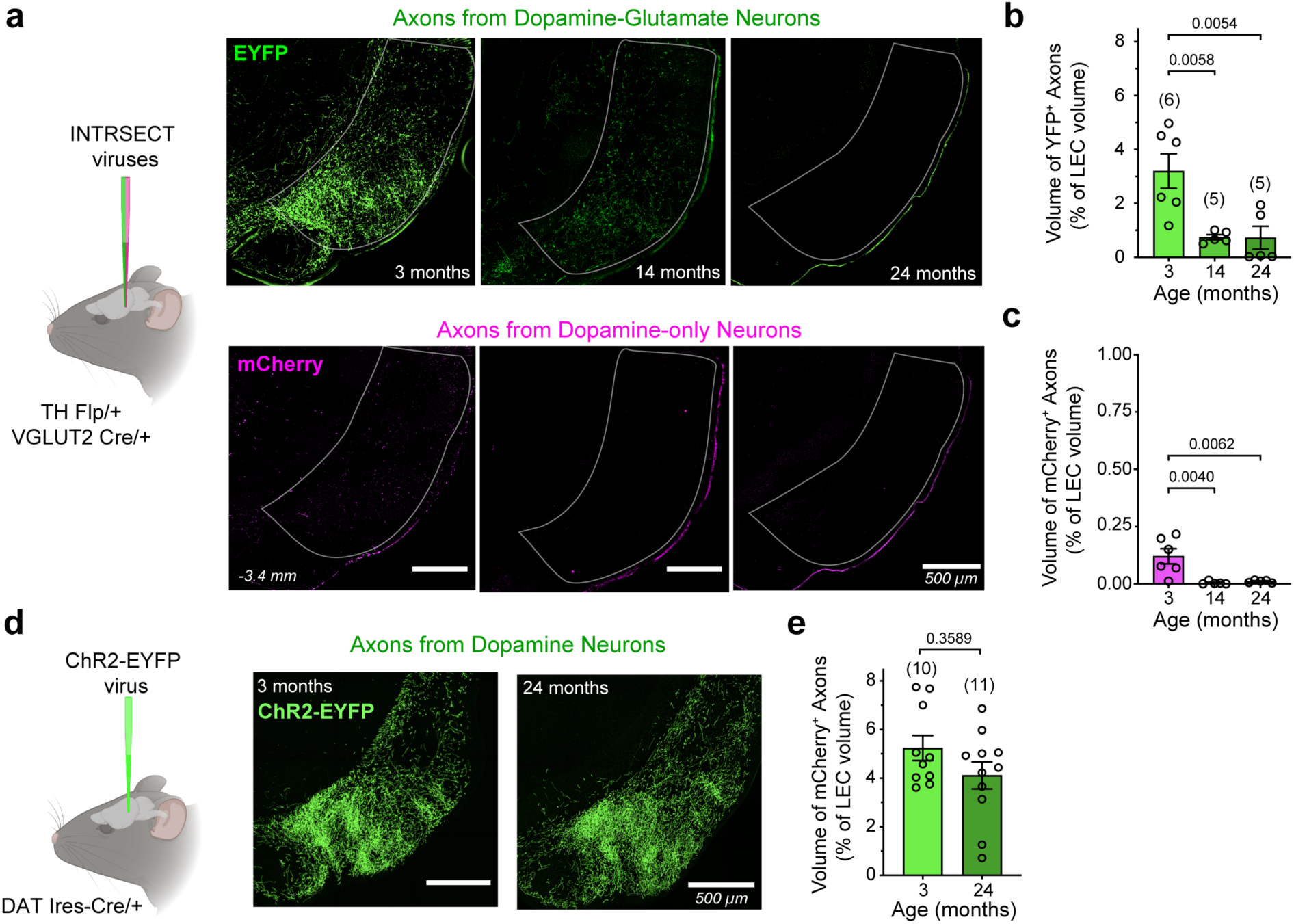
Distinct viral labeling strategies reveal an age-related reduction in INTRSECT EYFP+ DA-GLU axonal density in the LEC that is not attributable to axonal degeneration. **a,** Experimental schematic showing INTRSECT viruses injected into TH Flp/+ VGLUT2 Cre/+ mice (left) and representative images of EYFP+ DA-GLU axons (right top) and mCherry + DA-only axons (right bottom) in the outlined LEC at 3, 14, and 24 months. **b,** Volume of EYFP+ DA-GLU axons expressed as a percentage of total LEC volume showed a significant effect of age (Welch’s ANOVA: F(2, 6.36) = 6.48, p = 0.029; η² = 0.59). Dunnett’s multiple comparisons versus 3 months revealed significant reductions at 14 months (p = 0.0058) and 24 months (p = 0.0054). **c,** Volume of mCherry+ dopamine-only axons as a percent of LEC volume showed a significant effect of age (Welch’s ANOVA: F(2,8.12) = 6.68, p = 0.019; η² = 0.59). Dunnett’s multiple comparisons versus 3 months showed significant reductions at 14 months (p = 0.0040) and 24 months (p = 0.0062). **d,** Experimental schematic showing ChR2–EYFP virus injected into DAT-Ires-Cre/+ mice to label all dopaminergic axons (left) independently of co-transmission phenotype. Representative images of ChR2–EYFP+ axons in the LEC at 3 and 24 months (right). **e,** ChR2–EYFP+ axon volume as a percent of LEC volume did not differ between 3 and 24 months (Mann–Whitney, *U* = 68.5, *P* = 0.36; moderate effect size: *r* = 0.21). Error bars shown as mean ± SEM; numbers in parentheses above the bars indicate the number of mice.

We further investigated why INTRSECT labeling of DA-GLU neurons and their axons in the LEC declines with age. A recent study reported that TH and VGLUT2 RNA expression in the VTA decreases in aged mice, without evidence of cell loss ^65^. We therefore hypothesized that reduced expression of TH or VGLUT2 may underlie the diminished INTRSECT labeling, since this viral strategy depends on the activity of both promoters. To test this, we used an alternative viral strategy in which YFP is expressed under the DAT promoter, independent of TH or VGLUT2 expression status. An AAV-EF1a-DIO-ChR2(H134R)-EYFP virus was injected into the VTA of DAT-Ires-Cre/+ mice at 2 or 23 months of age to generate young (3-month-old, *n* = 10) and aged (24-month-old, *n* = 11) cohorts. With this approach, EYFP+ dopaminergic axons in the LEC were clearly observed in both groups (**Fig. 4d**), and quantification of terminal volume revealed no age-related differences (**Fig. 4e**). Thus, terminal density in the LEC is largely preserved at 24 months. Taken together, these results argue against cell loss or terminal degeneration and instead suggest that the reduction in INTRSECT labeling reflects decreased TH or VGLUT2 expression with age.

### Aging Effects on Dopamine Synthesis and Glutamate Vesicular Loading in DA-GLU axons in the LEC

We investigated whether EYFP-labeled dopaminergic axons in the DAT-Ires-Cre/+ mouse expressed the proteins TH and VGLUT2. To quantify TH expression, we performed volume rendering and generated surfaces of both TH and EYFP immunoreactivity (**Fig. 5a**). TH clusters overlapping by more than 65% with EYFP labeling were considered to be inside EYFP+ axons, and the volume of TH inside axons was calculated relative to the total volume of EYFP+ axons in the LEC. We quantified the superficial and deep layers of the LEC separately, as these layers are functionally distinct ^72–74^ and EYFP+ axons were denser in the superficial layer (**Extended Fig. 4a**). The expression of TH within EYFP+ axons was significantly reduced with age in both layers (**Fig. 5A, B, supplementary video 1 and 2**). However, TH immunoreactivity outside EYFP+ axons did not differ between young and aged mice (**Extended Fig. 4b**). Because noradrenergic neurons also innervate the LEC ^32^, it is plausible that the TH signal not associated with EYFP+ fibers reflects noradrenergic projections.

**Fig. 5.**
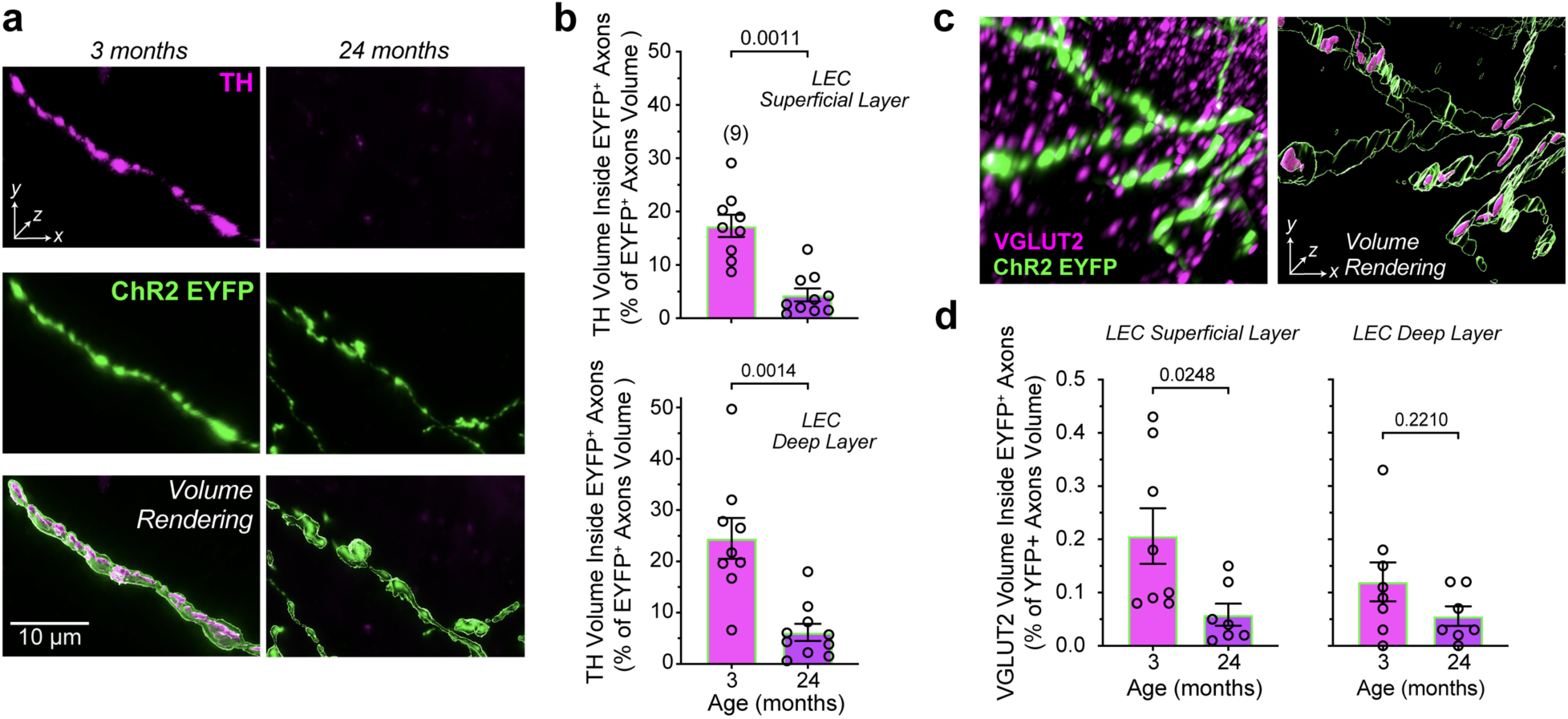
Age-related reduction of TH and VGLUT2 content in LEC-projecting DAergic terminals. **a**, Representative high-magnification images of axons in the LEC of 3- (left) and 24-month-old (right) DAT-Ires-Cre/+ mice expressing ChR2-EYFP in VTA dopamine neurons. Each column shows TH immunoreactivity (top, magenta), ChR2-EYFP signal (middle, green), and volume rendering (bottom) of merged signals from the same field of view. **b**, Quantification of TH signal volume within EYFP+ axons, expressed as a percentage of total EYFP+ axon volume, in superficial (top) and deep (bottom) layers of the LEC. Aged mice exhibited a significant reduction in TH content in both superficial and deep layers (superficial layer: Mann–Whitney U test, U = 8, r = 0.71; deep layer: Mann–Whitney U test, U = 9, r = 0.70; p-values above the brackets). **c,** Representative high-resolution image of VGLUT2 puncta (magenta) colocalized with EYFP-labeled axons (green) in the LEC; raw fluorescence image (left); 3D volume rendering of the same region (right). **d**, Quantification of VGLUT2 signal volume within EYFP+ axons, expressed as a percentage of total EYFP+ axon volume, in superficial (left) and deep (right) LEC. Aged mice showed a significant reduction in VGLUT2 content in the superficial layer (Mann–Whitney U test, U = 9, r = 0.58), with no significant difference in the deep layer (Mann–Whitney U test, U = 39, r = 0.32; p-values above the brackets). Scale bars, 10 µm.

We then assessed whether VGLUT2 expression is also reduced within EYFP-labeled axons. VGLUT2 immunolabeling appeared as densely clustered puncta in the LEC, reflecting expression in both DAergic axons and other glutamatergic inputs, with the latter providing the majority of VGLUT2. The overall volume of VGLUT2 puncta in the LEC did not change with aging (**Extended Fig. 4c)**. To quantify VGLUT2 inside YFP+ axons, we acquired images with a confocal microscope equipped with Airyscan super-resolution detection ^75^. Compared to conventional confocal microscopy, Airyscan employs a detector array with pixel reassignment and deconvolution, improving lateral resolution to ∼120 nm and enhancing signal-to-noise, thereby allowing a more reliable visualization of densely packed puncta. VGLUT2 puncta overlapping by more than 65% with YFP labeling were considered inside YFP+ axons (**Fig. 5c**). Using this conservative criterion, we found that in young mice, only ∼0.2 % of total VGLUT2 expression in the LEC was localized inside dopaminergic axons, confirming that the vast majority of VGLUT2 arises from non-dopaminergic inputs. As with TH, we analyzed superficial and deep layers separately, calculating the volume of VGLUT2 puncta as a percentage of total YFP+ axonal volume (**Fig. 5d**). In the superficial layer, VGLUT2 volume was significantly reduced in aged mice. In contrast, in the deep layer the difference between ages did not reach significance.

In conclusion, TH expression was robustly reduced within dopaminergic axons in the LEC across both superficial and deep layers. VGLUT2 expression declined with aging in a layer-specific manner, reaching significance only in the superficial LEC. Together, these findings indicate that DA-GLU neurons projecting to the LEC undergo marked age-related deterioration, with a substantial loss of dopamine synthesis capacity and a more moderate, regionally restricted decline in glutamate vesicular loading.

### Effects of Aging on Dopamine Release in the LEC

Decreases in TH expression are typically associated with reduced dopamine release ^76^; however, compensatory mechanisms are common in dopamine neurons and can preserve dopaminergic function ^77^. Thus, we evaluated the impact of TH reduction on dopamine release by optogenetically stimulating dopaminergic axons in the LEC with red-shifted ChRmine, and measuring dopamine dynamics with the green dopamine sensor GRAB_DA2h_ using fiber photometry (**Fig. 6a,b; Extended Fig. 5a**). We reasoned that if dopamine synthesis is compromised, the largest effects on release would emerge during repeated stimulation and at higher frequencies, when maintaining release relies strongly on ongoing dopamine synthesis. To first test for potential crosstalk between the red stimulation and green recording channels, we measured GRAB_DA2h_ signals in response to red light in the absence of ChRmine expression and observed no light-evoked signal, allowing us to continue with the designed experiment (**Extended Fig. 5b-e**). We stimulated dopaminergic axons with three consecutive 60-s bursts at four frequencies (5, 10, 20, and 40 Hz). The order of the 5, 10, and 20 Hz stimulation frequencies was randomly assigned, while 40 Hz stimulation was delivered at the end of the session to minimize the possibility that high-frequency–induced plasticity could affect responses at the other frequencies (**Fig. 6c, left**). For each frequency, bursts were delivered across three trials, responses were averaged, and mean traces were compared across frequencies (**Fig. 6c, right**).

**Figure 6.**
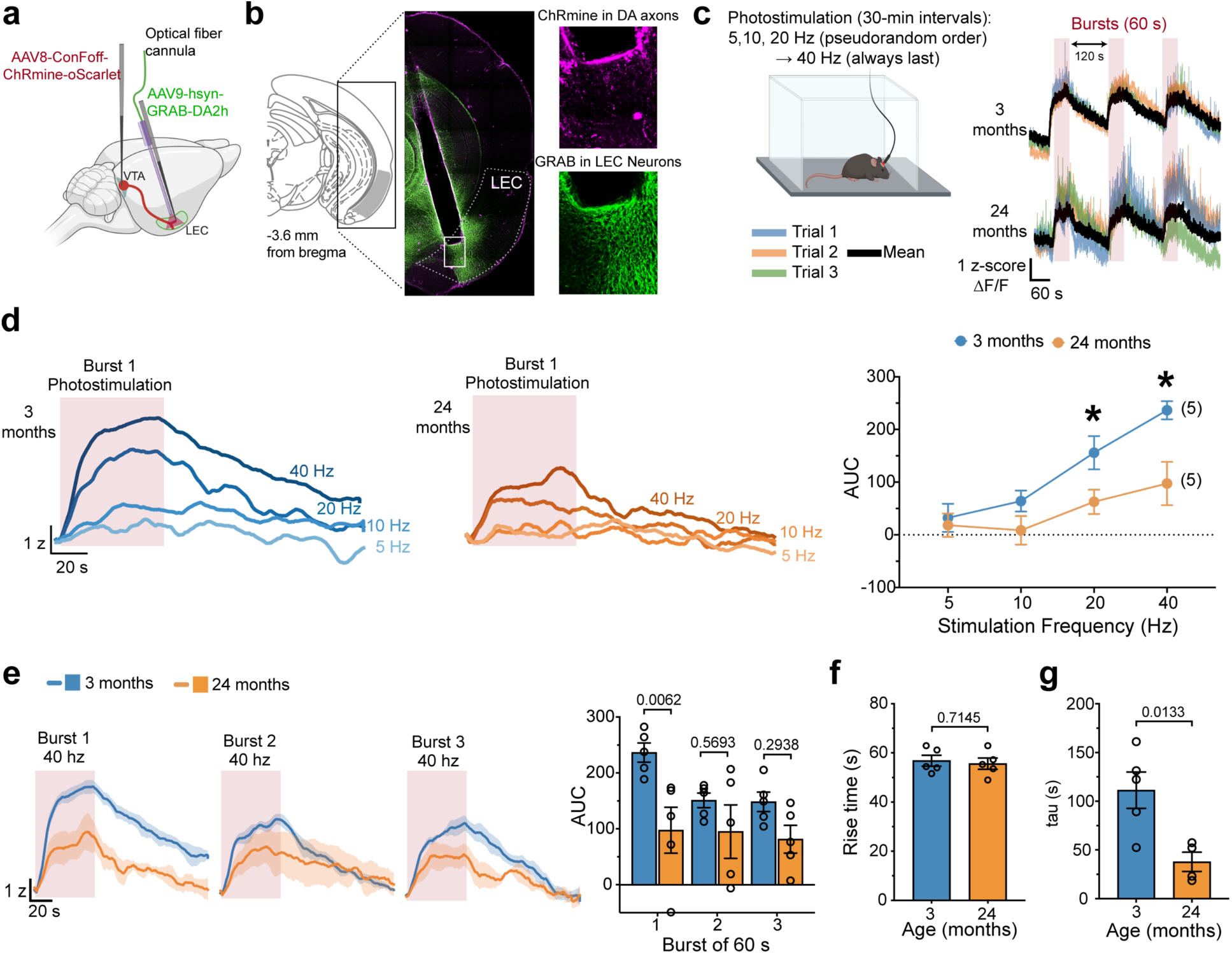
Aged mice show frequency-dependent decreases in evoked dopamine release. **a,** Schematic of viral delivery: AAV8-Con/Foff-ChRmine-oScarlet was injected into the VTA, and AAV9-hSyn-GRAB_DA2h_ was injected into the LEC at the site of optic fiber implantation for fiber photometry. **b**, Coronal section containing the LEC with representative photomicrographs showing optic fiber placement; higher-magnification insets beneath the probe illustrate ChRmine expression in dopaminergic terminals (magenta) and GRAB_DA2h_ expression (green) in LEC neurons. **c**, *In vivo* fiber photometry recordings were obtained in an open field during optogenetic stimulation. The order of 5, 10, and 20 Hz stimulation was randomized; 40 Hz stimulation was delivered at the end of the session. Representative raw traces in response to 40 Hz from two animals from different age groups (*right*). For each frequency, three bursts were delivered across three trials (blue, orange, and green traces), averaged (black trace) and compared across frequencies. **d**, Mean traces for the first burst (burst 1) at each stimulation frequency in young (blue) and aged (orange) mice. Right, area under the curve (AUC) as a function of age and frequency. Two-way repeated-measures ANOVA revealed a significant Age × Frequency interaction (Age × Frequency: F(2.278, 18.22)=4.612, P=0.0205), a main effect of frequency (F(2.278, 18.22)=28.68, P<0.0001), and a main effect of age (F(1,8)=5.700, P=0.0440). Planned comparisons at 20 and 40 Hz were performed using one-tailed unpaired Welch’s t-tests (Young > Aged) with Holm–Bonferroni correction across the two tests: young mice showed larger responses at 20 Hz (t(7.31)=2.39, P=0.0234; Holm-adjusted P=0.0237) and 40 Hz (t(5.37)=3.12, P=0.0119; Holm-adjusted P=0.0237), with large effect sizes (Hedges’ g=1.36 and 1.79, respectively). **e**, Mean traces for 40 Hz stimulation across three consecutive bursts (young, blue; aged, orange; shaded areas indicate SEM). Bar plot on the right shows a significant age effect on burst 1, but not on burst 2 or burst 3 (RM-ANOVA; significant Age x Burst interaction; Bonferroni-corrected p values shown above brackets). **f**, Rise time for the first 40 Hz burst, showing no effect of age. **g**, Decay time constant (τ) for the first 40 Hz burst, showing a significant reduction in aged mice (unpaired t-test, t=3.217, p value above the bracket).

We next examined the effect of stimulation frequency by comparing responses to the first burst. In both age groups, increasing stimulation frequency increased dopamine release, consistent with prior reports in other dopaminergic targets ^78, 79^ (**Fig. 6d**). However, responses differed by age only at higher frequencies, with no differences at 5 or 10 Hz but significantly reduced responses in aged mice at 20 and 40 Hz.

We next assessed how repeated stimulation influenced dopamine release across frequencies and ages by comparing responses to three 60-s bursts delivered at 120-s intervals (**Fig. 6c**, left). Age affected repeated-burst responses at 40 Hz (**Fig. 6e**), but not at 5, 10, or 20 Hz (**Extended Fig. 5f-h**). At 40 Hz, young mice showed the largest response during the first burst, followed by a decrease to a lower, stable level across subsequent bursts. In contrast, aged mice exhibited a reduced response during the first burst relative to young mice, with no further decline across consecutive bursts. Analysis of response kinetics during the first burst revealed no age effect on rise time (**Fig. 6f; Extended Fig. 5i**), but aged mice showed a significantly faster decay (reduced τ, **Fig. 6g**).

In summary, aged mice exhibited a frequency-dependent reduction in evoked dopamine release that emerged only at higher stimulation frequencies, consistent with reduced dopamine synthesis. The faster decay (shorter τ), often interpreted as enhanced dopamine clearance from the extracellular space ^80, 81^, may reflect compensatory increases in dopamine uptake that help stabilize evoked responses across stimulation frequencies.

## Discussion

The present study identifies a previously unrecognized, age-related decline in TH and VGLUT2 expression within a subpopulation of DA-GLU neurons projecting to the LEC, uncovering their selective vulnerability to aging.

Using intersectional viral strategies, we selectively labeled DA-GLU and DA-only neurons based on the presence or absence of glutamatergic and DAergic marker expression (VGLUT2 and TH, respectively) ^60, 63, 66^. These INTRSECT fluorescent reporters, administered at different ages, faithfully tracked endogenous TH and VGLUT2 expression and captured their age-related decline, providing a robust tool to monitor molecular changes within defined dopaminergic subpopulations. Using this approach, INTRSECT-driven EYFP labeling of DA-GLU neurons in the VTA and their axons in the LEC was markedly reduced in middle-aged and aged mice. The ∼40% reduction in VTA DA-GLU neuron density at middle age was accompanied by ∼75% reduction in labeled axons in the LEC, suggesting that LEC-projecting DA-GLU neurons are among the first VTA outputs to be affected during aging.

Despite the pronounced loss of INTRSECT-driven signal, there was no evidence of axonal degeneration, as LEC dopaminergic axons remained detectable in aged mice using an alternative DAT promoter-driven viral strategy. Instead, our data support decreased TH and VGLUT2 expression as the principal explanation for reduced INTRSECT labeling in the LEC in the absence of overt structural loss. Single-cell spatial transcriptomic studies in aged mice similarly suggest that VTA dopaminergic neuron abundance is preserved with age ^82^; and consistent with our findings, recent work further indicates that aging is associated with reduced TH and VGLUT2 transcript expression in VTA dopamine neurons without detectable cell loss ^65^. Interestingly, Buck et al., 2025 reported that these transcript reductions were not accompanied by decreased TH and VGLUT2 protein expression in the nucleus accumbens, a major target of DA-GLU neurons ^47, 56, 60^. In contrast, in our study TH protein levels and, to a lesser extent, VGLUT2 protein levels were reduced in DA-GLU axons in the LEC. Together, these findings suggest that DA-GLU mesolimbic projections may engage compensatory mechanisms that preserve protein expression despite transcript loss, whereas DA-GLU projections to the LEC show less compensation and increased vulnerability to aging.

The dopaminergic innervation of the LEC is denser in superficial layers (which provide major input to the hippocampus) than in deeper layers (which primarily relay hippocampal output). TH expression was reduced to a similar extent in both compartments, whereas VGLUT2 expression was significantly reduced only in superficial layers. These convergent effects in superficial layers raise the possibility that the LEC-to-hippocampus input pathway is especially vulnerable to aging-related DA–GLU decline.

Our fiber photometry data show that the decrease in TH expression has functional consequences and compromises dopamine release in a frequency-dependent manner. As predicted, the greatest deficits emerged during high-frequency firing when rapid synthesis is required to sustain release. Dopamine neurons typically exhibit pacemaker-like firing at ∼4 Hz but transiently increase their activity in bursts of 20–100 Hz to signal salient events in the environment, such as novel stimuli or predictive cues ^27, 29, 83^. Recordings from DA-GLU neurons have shown that the burst firing of these neurons encodes the salience of a stimulus independently of its valence, whether rewarding or aversive ^84^. This phasic mode of salience signaling may be particularly vulnerable to the age-related reduction in TH expression.

Aging-related preservation of VTA cell number stands in contrast to Alzheimer’s disease patients and transgenic mouse models, which show dopaminergic neurodegeneration in the VTA ^85, 86^, particularly within its middle portion, where DA-GLU neurons are concentrated. Thus, the molecular alterations we identify in DA-GLU neurons projecting to the LEC during normal aging may represent an early vulnerability state that precedes the overt neurodegenerative phenotypes observed in Alzheimer’s disease. In post-mortem tissue from individuals with Parkinson’s disease, there is evidence that melanized dopamine neurons in the substantia nigra lose TH expression before undergoing neurodegeneration ^87^. Recent evidence suggests that functional impairments in LEC-projecting dopamine neurons also occur in Alzheimer’s mouse models ^88^. This study shows that dopamine-related activity in the LEC was reduced during an associative learning task, with blunted responses to novel odors, leading to associative memory deficits. Because these effects were observed in young AD mice, when dopaminergic neurodegeneration is limited, the deficits likely reflect, at least in part, impaired recruitment of LEC-projecting dopamine neurons by salient environmental cues. The rescue with L-DOPA further suggests that reduced dopamine synthesis contributes to this phenotype, paralleling the dopamine-related functional impairments we observe in aged mice.

Taken together, these findings position DA-GLU projections to the LEC as an early “weak link” in the mesocorticolimbic system, one that could cascade into a disproportionate disruption of LEC-hippocampal-dependent memory and salience processing.

## Methods

### Experimental Animals

All procedures were conducted in accordance with National Institutes of Health *Guide for the Care and Use of Laboratory Animals* and approved by the Institutional Animal Care and Use Committee of the City University of New York (CUNY) Advanced Science Research Center (ASRC). Mice were housed under standard barrier conditions with a reverse 12-hour light/dark cycle (light off from 11 am to 11 pm) and ad libitum access to food and water. Male and female mice were used for all experiments. Two transgenic lines were utilized: (1) TH-Flp/+ ; VGLUT2-Cre/+ mice for INTRSECT viral experiments and (2) DAT-Ires-Cre/+ mice for ChR2-YFP labeling. Animals were aged to 2-3 months (young), 13-14 months (middle), or 23-24 months (old).

### Stereotactic Surgery

#### INTRSECT Viral Strategy

To selectively label DA-GLU and DA-only VTA neurons, TH-Flp/+ ;VGLUT2-Cre/+ mice were injected with a mixture of AAV8-EF1a-Con/Fon-EYFP-WPRE and AAV8-EF1a -Coff/Fon-mCherry-WPRE ^66^. The viruses were kindly donated by Dr. Deisseroth’s lab. Injections were performed at 2, 13, or 23 months of age to allow one month of viral expression prior to tissue collection at 3, 14, or 24 months (n = 9, 7, and 5, respectively). Mice were anesthetized with isoflurane (3% induction, 2% maintenance), and 1 μL of viral solution (1.5 × 10^12^ vg/mL) was pressure-injected into the unilateral medial VTA and lateral VTA/SNc (Nanoject III, Drummond Scientific). Stereotaxic coordinates (relative to bregma) were adjusted by bodyweight: medial VTA: AP –3.0 to –3.4 mm, ML ±0.5 mm, DV –4.1 to –4.5 mm; lateral VTA/SNc: AP –3.0 to –3.4 mm, ML -1.3 mm, DV –4.3 mm. The pipette remained in place for 5 minutes to minimize backflow.

#### ChR2-YFP Strategy

To label dopaminergic axons independent of TH or VGLUT2 promoter activity, DAT-Ires-Cre mice were injected bilaterally in the VTA with AAV5-EF1a-DIO-ChR2(H134R)-EYFP (UNC Lot # AV4313-2B; 1.5 × 10^12^ vg/mL) at either 2 months (n = 10) or 23 months (n = 11). Coordinates for the medial VTA and procedure matched the INTRSECT protocol above.

#### Fiber Optic Implantation Combined with Viral Injections

To express a red-shifted opsin, DAT-IRES-Cre mice received unilateral injections of AAV8-nEF1-Con/Foff 2.0-ChRmine-oScarlet (Addgene #137161; 2.16 x 10^12^ vg/mL) into the VTA using the medial VTA coordinates described above. Mice also received bilateral injections of AAV9-hsyn-GRAB_DA2h (Addgene #140554; 1.88 x 10^12^ vg/mL) into the LEC (AP –3.4 mm, ML -0.5 mm, DV –3.8 to –3.9 mm, angle 8°). A fiber optic cannula (black ceramic ferrule, 1.25 mm ferrule diameter; 200 µm core; NA = 0.37; length = 5.0 mm; Neurophotometrics) was implanted into the LEC at the same coordinates used for the GRAB_DA2h_ injection.

### Immunohistochemistry

#### INTRSECT-Labeled Brains

Immunohistochemistry was performed as previously described ^61^. Mice were anesthetized (ketamine/xylazine) and perfused with PBS followed by 4% paraformaldehyde (PFA). Brains were post-fixed and cut into 50 μm coronal sections in a vibratome (MicroSlicer DTK-1000N). Sections underwent sequential washes in 1x PBS (3 x 10 minutes), followed by incubation in 0.1 M glycine, washed in 1x PBS again (3 x 10 minutes), then were blocked in 10% normal goat serum (NGS) with 0.1% Triton X-100 and incubated with primary antibodies: anti-mCherry (rat, 1:5000; Thermo Fisher Scientific Cat# M11217, RRID:AB_2536611), anti-GFP (chicken, 1:5000; Thermo Fisher Scientific Cat# A10262, RRID:AB_2534023) and either anti-TH (rabbit, 1:5000; Thermo Fisher Scientific Cat# OPA1-04050, RRID:AB_325653) or anti-VGLUT2 (rabbit, 1:250–1:5000; Synaptic Systems #135403; RRID:AB_887883). Secondary antibodies anti-rat Alexa Fluor 568 (goat, 1:200; Thermo Fisher Scientific Cat# A-11077, RRID:AB_2534121) anti-chicken Alexa Fluor 488 (goat, 1:200; Thermo Fisher Scientific Cat# A32931, RRID:AB_2762843) and anti-rabbit Alexa Fluor 647 (goat, 1:200 to 1:400; Thermo Fisher Scientific Cat# A32733, RRID:AB_2633282) diluted in 0.02% PBS-T. Sections underwent additional washes in 1x PBS (3 x 10 minutes) and mounted and coverslipped with ProLong Gold Antifade mounting medium (Thermo Fisher Scientific).

#### ChR2-YFP Labeled Brains

Sections were processed similarly as the INTRSECT-labeled brains but blocked with an adapted aging-specific solution prepared using PGBA, 10% NGS, and 0.5% Triton X-100. Primary antibodies for TH quantification: anti-GFP (rabbit, 1:5000; Abcam Cat# ab6556, RRID:AB_305564) and anti-TH (chicken, 1:5000; Aves Labs Cat# TYH, RRID:AB_10013440). Primary antibodies for VGLUT2 quantification: anti-GFP (chicken, 1:5000; Thermo Fisher Scientific Cat# A10262, RRID:AB_2534023) and anti-VGLUT2 (rabbit, 1:1000: Synaptic Systems Cat# 135403, RRID:AB_887883). Secondary antibodies for TH quantification: anti-rabbit Alexa Fluor Plus 647 (goat, 1:400; Thermo Fisher Scientific Cat# A32733, RRID:AB_2633282) and anti-chicken Alexa Fluor 568 (goat, 1:400; Thermo Fisher Scientific Cat# A-1104, RRID:AB_2534098). Secondary antibodies for VGLUT2 quantification: anti-chicken Alexa Fluor Plus 647 (goat, 1:400; Thermo Fisher Scientific Cat# A32933, RRID:AB_2762845) and anti-rabbit Alexa Fluor 532 (goat, 1:400; Thermo Fisher Scientific Cat# A-11009, RRID:AB_2534076). Secondary antibodies then underwent a wash in deionized water (5 minutes). Lipofuscin autofluorescence was quenched using cupric sulfate (CuSO) in ammonium acetate (50 mM, pH 5.0; 15 minutes). A subsequent wash in deionized water (5 minutes) followed, after which sections underwent additional washes in 1x PBS (3 x 10 minutes) and mounted and coverslipped with ProLong Gold Antifade mounting medium (Thermo Fisher Scientific).

### In Situ Hybridization (RNAscope) Combined with Immunohistochemistry

*In situ* hybridization was combined with immunohistochemistry using RNAscope reagents (Advanced Cell Diagnostics; ACD Bio) as previously described ^30^ and was performed by the Epigenetics Core Facility at the CUNY Advanced Science Research Center (ASRC). Tissue sections were fixed in chilled 4% paraformaldehyde for 15 min at 4 °C, dehydrated through graded ethanol, and air-dried for 5 min. Endogenous peroxidase activity was quenched with hydrogen peroxide for 10 min, followed by antigen retrieval for 5 min in boiling buffer. Sections were then processed for IHC with anti-GFP (chicken, 1:1000) and anti-mCherry (rat, 1:1000), followed by protease digestion for 30 min at 40 °C. For RNAscope, the vesicular glutamate transporter 2 probe (Slc17a6/VGLUT2; ACD Bio, #1170921-C2) was hybridized for 2 h at 40 °C in a humidity-controlled oven (HybEZ II, ACD Bio). Signal amplification was performed using RNAscope AMP reagents (ACD Bio), and probe signal was visualized using probe-specific horseradish peroxidase–based detection with Opal 690 dye (PerkinElmer; 1:1500). Slides were subsequently incubated with secondary antibodies (anti-chicken Alexa Fluor 488, 1:400; anti-rat Alexa Fluor 568, 1:400), counterstained with DAPI, and coverslipped with ProLong Gold Antifade mounting medium (Thermo Fisher Scientific).

### Image Acquisition and Quantification

#### INTRSECT quantification in the VTA

Some images were acquired on a Leica DM6 B epifluorescence microscope, and others on a Zeiss LSM 880 confocal microscope. Because quantification did not differ across imaging modalities, data from both systems were pooled for analysis. Neuronal subtypes within the VTA were quantified in ImageJ. For each z-stack, a z-projection was generated to create a reference image for counting. Raw optical sections were viewed in parallel to confirm signal localization within somata and to avoid misclassification due to projection artifacts. Manual cell counts were performed using the ImageJ Multi-point Tool. Each neuronal phenotype (and each of the seven combinatorial marker categories) was assigned a unique counter index: TH-only, mCherry-only, EYFP-only, TH+mCherry, TH+EYFP, TH+mCherry+EYFP, and mCherry+EYFP. Cells were classified using predefined inclusion criteria based on marker colocalization and somatic morphology. Each counted cell was marked on the z-projected image and then verified against the corresponding raw z-stack to ensure accurate attribution of fluorescence within individual cell bodies. Cell counts and x–y coordinates were exported from ImageJ and compiled in Microsoft Excel for organization and downstream analysis. Total counts per subtype and per section were calculated from the exported files prior to statistical analysis. The exported x–y coordinates were used to generate 2D spatial plots of cell distributions.

#### INTRSECT-labeling and VGLUT2 RNAscope Quantification in the VTA

Images were acquired on a Zeiss LSM 880 confocal microscope. All analyses were performed in Imaris (Bitplane/Oxford). The VTA was defined in Imaris by manual surface delineation. Boundaries were traced on the first optical section and propagated across the full z-stack. The VTA surface was then used to mask the relevant fluorescence channels to restrict analysis to the region of interest (ROI) (mCherry and EYFP channels). Cell bodies were segmented from the masked neuronal fluorescence channels as 3D surfaces using Imaris surface creation (smoothing enabled; surface detail = 2.0 µm), with intensity thresholding and background subtraction applied to separate signal from background. Segmented neuronal surfaces were manually curated to remove artifacts, merge fragmented objects, and split incorrectly merged cells. VGLUT2 mRNA transcripts were quantified using Imaris spot detection on the masked VGLUT2 transcript channel. The spot (XY) diameter was determined empirically by measuring puncta in slice view and then held constant across animals within the VTA. To quantify VGLUT2 puncta per neuron, the Imaris Cell to Vesicle function was used. Pre-segmented neuronal surfaces (mCherry+ and EYFP+ cells) were imported as “cell” objects, and detected VGLUT2 spots were imported as “vesicle” objects. Spots were classified as intracellular for a given neuron when the distance from the spot to that neuron’s surface was zero; spots with nonzero distance were considered extracellular relative to that neuron. Per-cell transcript abundance was extracted as **“**Cell number of vesicles**,”** representing the number of VGLUT2 spots assigned to each neuronal surface.

#### INTRSECT-labeling and ChR2-EYFP Axonal Quantification in the LEC

Images were acquired on a Leica DM6 B epifluorescence microscope. All analyses were performed in Imaris (Bitplane/Oxford). In Imaris, the LEC ROI was manually delineated using anatomical boundaries and masks were generated on the relevant fluorescence channels (EYFP and/or mCherry) to restrict subsequent analysis to the selected signal. Within the selected LEC ROI, labeled axons were segmented using Imaris surface reconstruction without smoothing or background subtraction. Total axonal volume for each region was obtained from the Imaris surface statistics and used as the measure of axonal signal within the LEC and normalized by the total volume of the LEC.

#### TH Inside ChR2-EYFP+ Axons Quantification in the LEC

Images were acquired on a Leica DM6 B epifluorescence microscope and analyzed in Imaris (Bitplane/Oxford). For each section, regions of interest corresponding to superficial and deep LEC layers were defined using anatomical landmarks. The analysis window was held constant for each ROI (x–y dimensions fixed at 1000 × 1000 pixels, with a z-stack of 15 optical planes). Surface segmentation was restricted to the appropriate ROI to generate superficial and deep EYFP surfaces (EYFP+ axons) and superficial and deep TH surfaces (TH signal in axons). Colocalization was assessed by filtering TH surfaces based on their overlap with the corresponding EYFP surface using an overlapped volume ratio metric; TH signal was classified as within EYFP+ axons when the overlap ratio was ≥0.65. For each superficial and deep LEC ROI, the total TH volume classified as inside versus outside EYFP+ axons was extracted and recorded.

#### VGLUT2 Inside ChR2-EYFP+ Axons Quantification in the LEC

Images were acquired on a Zeiss LSM 880 confocal microscope equipped with Airyscan. Quantification followed the same analysis pipeline described for “TH Inside ChR2-EYFP+ Axon Quantification in the LEC”. Surfaces representing EYFP+ axons and VGLUT2+ axonal terminals were generated and filtered using the same overlap criteria of ≥0.65. For each superficial and deep LEC ROI, the total volume of VGLUT2 signal classified as within EYFP+ axons was extracted and recorded.

### Dual Color Fiber Photometry with Optogenetic Stimulation

#### ChRmine Stimulation and GRAB_DA2h_ Recordings

Fiber photometry recordings were acquired using a Neurophotometrics FP3002 system. Optogenetic stimulation was delivered through the LEC recording fiber optic cannula using a 635 nm laser (3 mW at the patch-cord tip) at 5, 10, 20, and 40 Hz (5 ms pulse width). For each frequency, stimulation consisted of a 60-s burst delivered three times with an inter-trial interval (ITI) of 120 s. The 5-, 10-, and 20 Hz conditions were presented in randomized order and separated by 30 minutes; the 40 Hz condition was always delivered last to avoid potential carryover effects of high-frequency stimulation on responses measured at lower frequencies. Simultaneously, GRAB_DA2h_ fluorescence was excited with a 470 nm LED (20 µW at the patch-cord tip). Emitted fluorescence was collected through a green bandpass filter and sampled at 30 Hz.

Analysis of the dopamine sensor signal was performed using custom-written Python scripts (MetaCell). We used the 470 nm sensor signal itself to generate a fitted baseline for photobleaching correction. Specifically, the 470 nm channel was detrended by fitting a biexponential curve across the entire session, and the raw 470 nm signal was then divided by this scaled fitted baseline to yield normalized fluorescence values (dF). Signals were smoothed using a 10 s rolling window, and peri-event time histograms were generated on a trial-by-trial basis from −2 to +180 s relative to stimulation onset. Baseline was defined as the mean signal from −2 to 0 s. Responses were expressed as z-scores. To quantify stimulation-evoked dopamine release, area under the curve (AUC) was calculated from 0–120 s relative to stimulation onset. For decay kinetics (τ_decay), the maximum baseline-corrected response was identified within a post-onset peak-search window of 0–65 s to obtain peak amplitude and its time (t_peak; used as the rise-time parameter). The decay phase was fit from t_peak to 180 s with a single-exponential model, y(t) = A·exp(−(t − t_peak)/τ_decay), using nonlinear least-squares minimization with constraints A > 0.5 and 0.05 ≤ τ_decay ≤ 500 s. Goodness-of-fit was assessed using the coefficient of determination (R²) computed over the fit window. For traces with poor fits (R² < 0.7), an alternative approach was used to prevent the offset term from dominating the fit: the offset was estimated as the mean signal from 95–100 s, and the decay segment (65–100 s) was offset-corrected and fit with a no-offset exponential model.

### Statistical Analyses

All statistical tests are reported in the figure legends. Statistical analyses were performed using GraphPad Prism or RStudio; with α set at 0.05. Data distributions were assessed using the Shapiro–Wilk test to determine whether parametric or nonparametric tests were appropriate. Between-group comparisons were analyzed using one-way ANOVA, two-way ANOVA, or repeated-measures ANOVA, followed by Dunnett’s or Bonferroni post hoc tests. Welch’s t-test was used to compare the means of two groups when the assumption of equal variances was not met. For non-normally distributed data, Kruskal–Wallis and Mann–Whitney U tests were used with Dunn’s post hoc correction when appropriate. Effect sizes are reported as partial eta squared (ηp²), Cohen’s d, or rank-biserial r, depending on the analysis.

To resolve the spatial distribution of neurons in the VTA, we labeled two points in each image, for each age category, that are indicative of the mid-line. Using these points, a straight line was drawn, and the distance of each cell type of interest from this midline was estimated. This is reflective of the medial-lateral distribution of the neurons. Owing to variation in size between ages and sexes, we normalized the distances from the midline for each image to the longest distance among all TH+ neurons. This enabled us to spatially compare the distributions of DA-only (mCherry+), DA-GLU (EFYP+), and TH+ neurons within, and between ages. For each image and each neuronal group, the normalized distances were used to compute a histogram. This allowed us to count the number of neurons at specific distances from the midline. For each age group, all this information was then sorted into a two-dimensional spatial map for each replicate animal, with bregma along one axis, medial-lateral distance on the other axis, and the counts of cells of interest for each combination of bregma and medial-lateral distance on the color axis. The spatial maps were then averaged over replicate animals for each age group, and the averaged spatial maps are represented as heatmaps. Difference heatmaps are generated by subtracting the averaged spatial maps. To compare the differences between spatial maps of different age groups, we carried out multiple one-sampled t-tests. For each bregma and medial-lateral distance, the replicate counts for 14- and 24-month-old animals were tested for statistical significance using a one-tailed t-test against the mean of the 3-month-old group for each cell (xy coordinate) in the map. Owing to the large number of t-tests, we then corrected the p-values using a Benjamini-Hochberg correction to adjust the FDR to 5%. Only the corrected p-values<0.05 were considered significant and are represented in the p-value heatmaps. All analyses, plots, and statistical comparisons were carried out using custom R scripts.

## Acknowledgments

This work was supported by the National Institutes of Health, through the National Institute of Mental Health under award MH123926 and the National Institute on Aging under award AG086477 (SM). JN was supported by the National Institute of General Medical Sciences of the National Institutes of Health through the Maximizing Access to Research Careers (MARC) Program under award T34GM007639, and by the National Institutes of Health under award R25GM065096. The authors acknowledge the Epigenetics Core Facility of the CUNY Advanced Science Research Center for instrument use and scientific and technical assistance with the *in situ* hybridization (RNAscope) experiment. Figures were created with BioRender.com.

## Ethics Declaration

The authors declare no competing interests.

---

**Extended Fig. 1.**
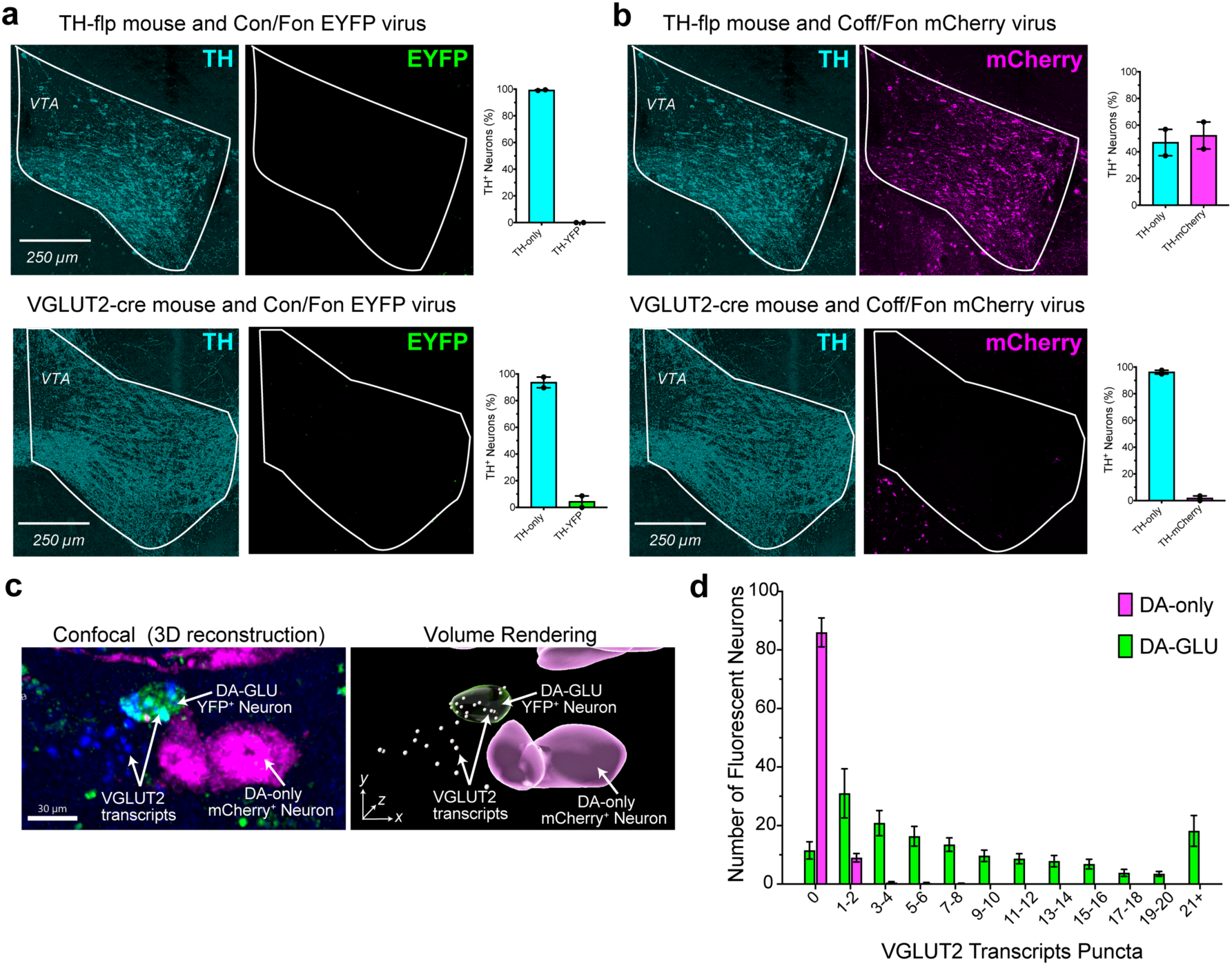
Validation of recombinase dependence for INTRSECT viral expression. **a**, Representative photomicrographs showing TH and EYFP immunoreactivity in TH-Flp (top) and VGLUT2-Cre (bottom) mice injected with an INTRSECT Con/Fon-EYFP virus. Quantification (right) shows no EYFP expression, confirming dual recombinase dependence. **b**, Representative photomicrographs showing TH and mCherry immunoreactivity in TH-Flp (top) and VGLUT2-Cre (bottom) mice injected with an INTRSECT Coff/Fon-mCherry virus. Quantification (right) shows robust mCherry expression in TH-Flp mice but no expression in VGLUT2-Cre mice, consistent with Flp-dependent activation and Cre-dependent suppression. **c**, Confocal images (left) and 3D renderings (right) showing VGLUT2 (Slc17a6) transcript puncta (blue) within EYFP+ neurons (green; DA–GLU) and mCherry+ neurons (magenta; DA-only). **d**, Quantification of VGLUT2 transcript puncta per neuron in EYFP+ versus mCherry+ populations shows that VGLUT2 (a glutamatergic marker) is enriched in EYFP+ neurons, whereas mCherry+ neurons exhibit little to no VGLUT2 signal, with only a few cells containing a single punctum.

**Extended Fig. 2.**
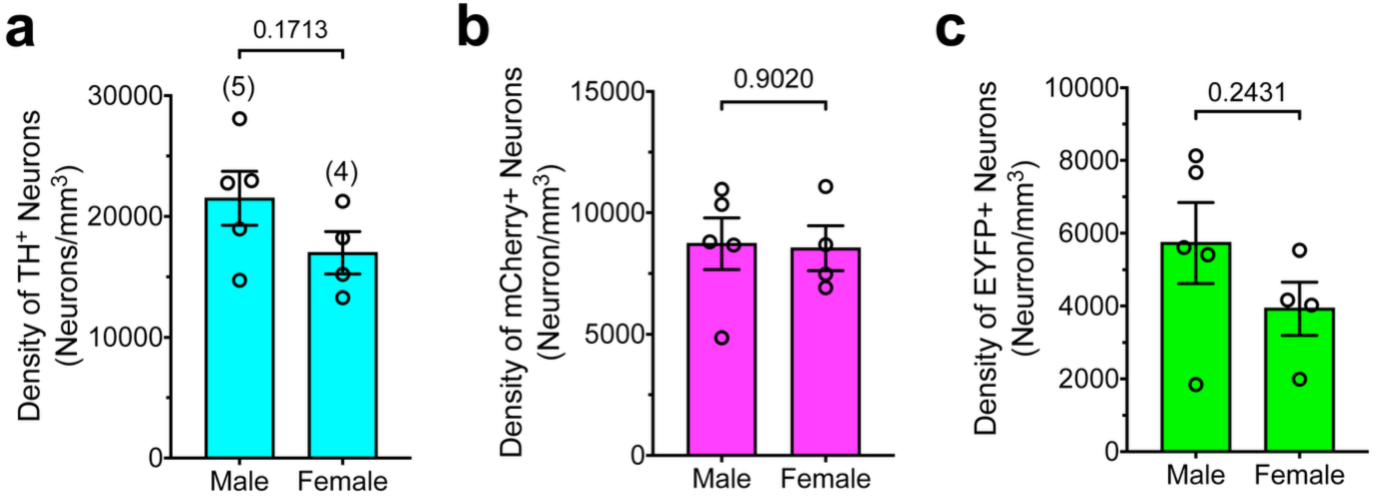
No sex differences in dopaminergic subpopulations density in the VTA of young mice. Bar graphs comparing the density (neurons/mm³) of TH+ neurons (**a**), mCherry+ neurons (**b**), and EYFP+ neurons (**c**) in male versus female young mice. Each bar represents group mean ± SEM. P-values are displayed on the plots, indicating no statistically significant differences between sexes for any neuronal population shown.

**Extended Fig. 3.**
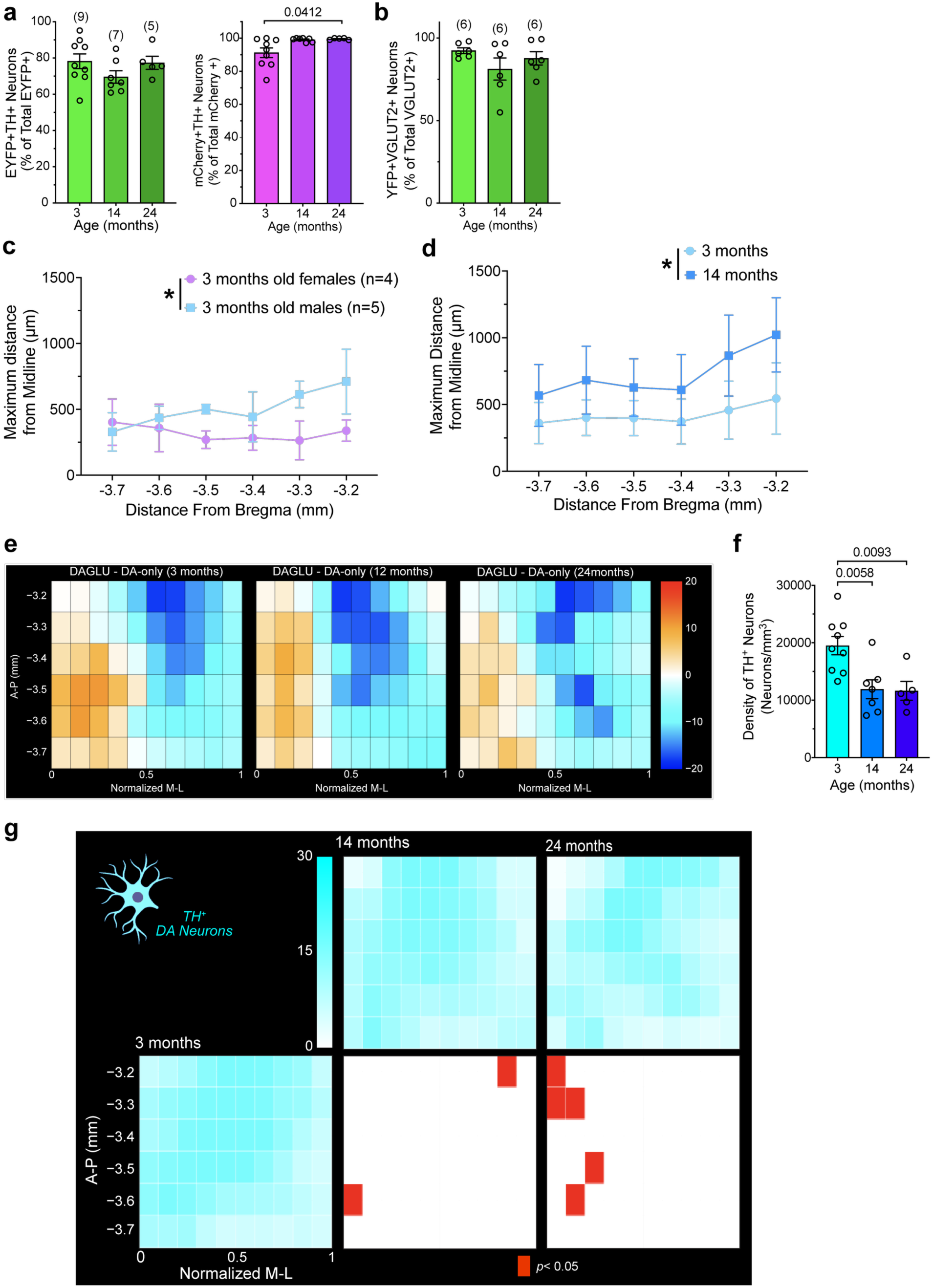
Validation of INTRSECT labeling in aged mice and age-related distribution of dopaminergic subpopulations in the VTA. **a,** Specificity of INTRSECT-labeled neurons for TH+ dopamine neurons at 3, 14, and 24 months. Left, percentage of EYFP+TH+ neurons among all EYFP+ cells; right, percentage of mCherry+TH+ neurons among all mCherry+ cells. Viral specificity was maintained with age, with a modest increase in TH co-localization in the mCherry+ population at 24 months (EYFP: no age effect, one-way ANOVA, F(2,18)=0.9193; mCherry: Kruskal–Wallis, H=6.992, Dunn’s post hoc; significant adjusted p-values indicated). Numbers above bars indicate mice. **b**, Validation of glutamatergic identity of EYFP+ neurons using EYFP immunoreactivity combined with in situ hybridization for VGLUT2 (*Slc17a6*). Plot shows the percentage of EYFP+VGLUT2+ neurons among all VGLUT2+ neurons at 3, 14, and 24 months. Each dot represents a section (4 mice per age, 6 sections per mouse). **e**, 2D VTA density maps showing the difference in spatial distribution between EYFP+ (DA–GLU) and mCherry+ (DA-only) cells across anterior–posterior (A–P) and normalized medial–lateral (M–L) coordinates. Color indicates the DA-GLU – DA-only difference per bin: white denotes no difference, warm colors indicate bins enriched in DA-GLU neurons, and cool colors indicate bins enriched in DA-only neurons. **f**, Bar graph showing TH+ neuron density (neurons/mm³) across age groups (one-way ANOVA main effect of age: *F*_(2,18)_ = 7.91, p=0.0034, large effect size: ηp²=0.47; Dunnett’s post hoc test vs 3 months; exact p values shown above brackets). **g**. 2D VTA density map showing the distribution of TH+ neurons across anterior–posterior (A–P) and normalized medial–lateral (M–L) coordinates. Red tiles indicate bins that differed from the 3-month group (one-tailed t-test vs the 3-month mean), with p-values Benjamini–Hochberg corrected to control the false discovery rate (FDR) at 5%; only adjusted p < 0.05 bins are shown.

**Extended Fig. 4.**
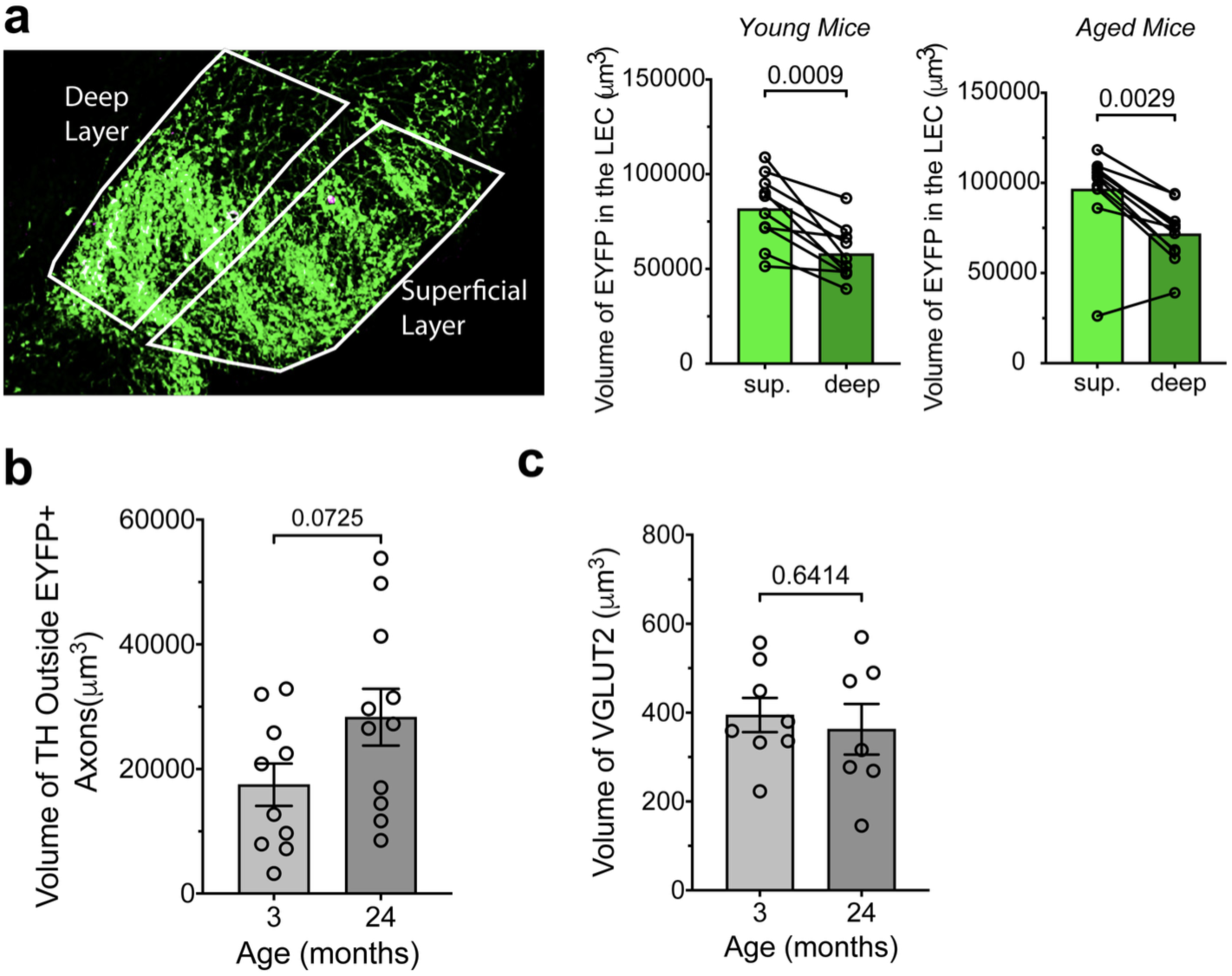
Layer- and age-dependent analysis of EYFP, TH, and VGLUT2 volume in the LEC. **a**, Left, representative LEC image illustrating the delineation of superficial and deep layers for volumetric analysis of ChR2–EYFP labeled axons in DAT-IRES-Cre mice. Bar plots show total EYFP+ axonal volume (µm³) in superficial versus deep layers at 3 (*left*) and 24 months (*right*). EYFP+ volume was greater in superficial than deep layers at both ages (3 months: paired t-test, *t*(9)=4.882, *P*=0.0009; 24 months: Wilcoxon matched-pairs signed-rank test, *W* = −62, *P*=0.0029). **b**, TH+ axonal volume (µm³) outside EYFP+ axons in the LEC at 3 and 24 months; no age difference (unpaired t-test, *t*(19)=1.876, *P*=0.0761). **c**, Total VGLUT2+ terminal volume (µm³) in the LEC at 3 and 24 months; no age difference (unpaired t-test, *t*(13)=0.4768, *P* = 0.6414). **d**, VGLUT2+ volume (µm³) in superficial and deep layers at 3 and 24 months; no age effect (unpaired t-test, *t*(13)=0.4768, *P* = 0.6414). Each dot represents an animal; bars show mean ± SEM.

**Extended Fig. 5.**
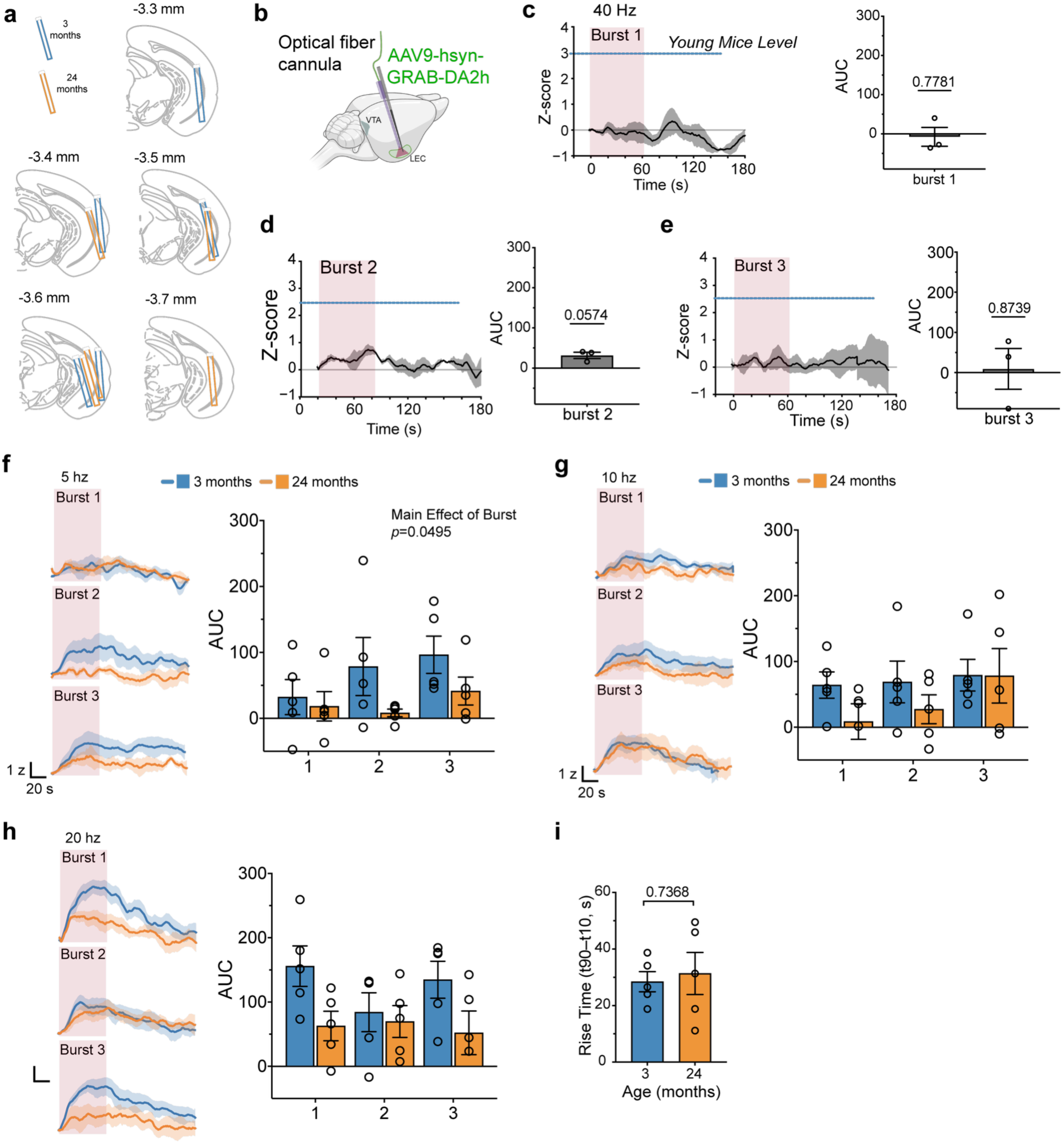
Validation of combined optogenetics and fiber photometry and dopamine responses to 5–20 Hz stimulation. **a,** Schematic of optic fiber cannula placement in young and aged mice. **b**, Schematic of injection of GRAB_DA2h_ virus and cannula placement for control experiments testing the effect of red-light stimulation on GRAB_DA_ signals. **c–e**, Peri-event histograms showing mean GRAB_DA_ responses to 40 Hz stimulation delivered as three consecutive 60-s bursts (n = 3 mice; 2 young, 1 aged; shaded area, ± SEM). Right, quantification of red-light effects on GRAB_DA_ signals for bursts 1–3; one-sample t-tests showed no significant effect (burst 1: t(2)=0.3218, P=0.7781; burst 2: t(2)=3.991, P=0.0574; burst 3: t(2)=0.1797, P=0.8739). **f**, Mean traces (young, blue; aged, orange; shaded area, ± SEM) and quantification for 5 Hz stimulation across three consecutive bursts. Two-way repeated-measures ANOVA showed no main effect of age and no Age × Burst interaction, but a main effect of burst (Age × Burst: F(2,16)=1.607, P=0.2312; Burst: F(2,16)=3.648, P=0.0495; Age: F(1,8)=1.935, P=0.2016). **g**, Same as f for 10 Hz stimulation; no significant effects (Age × Burst: F(2,16)=0.6000, P=0.5607; Burst: F(2,16)=1.428, P=0.2826; Age: F(1,8)=1.432, P=0.2657). **h**, Same as f for 20 Hz stimulation; no significant effects (Age × Burst: F(2,16)=2.824, P=0.0890; Burst: F(2,16)=1.620, P=0.2287; Age: F(1,8)=3.222, P=0.1104). **i**, Rise time (time to 90% peak minus time to 10% peak) did not differ by age (unpaired t-test, t(8)=0.3528, P=0.7333).

