## Supplemental for "Selective Vulnerability of Dopamine-Glutamate Neurons in Aging Weakens Entorhinal Dopamine Signaling"

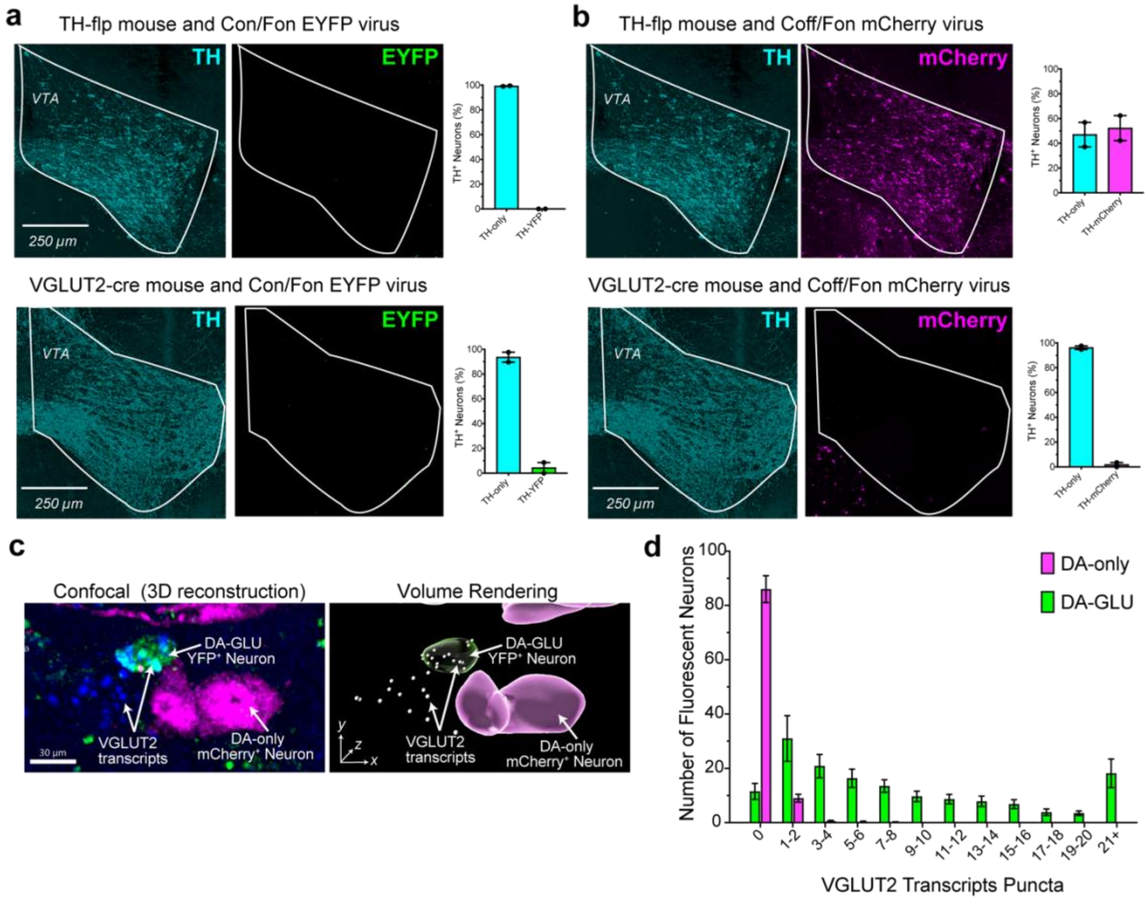


**Extended Fig. 1 | Validation of recombinase dependence for INTRSECT viral expression**. **a**, Representative photomicrographs showing TH and EYFP immunoreactivity in TH-Flp (top) and VGLUT2-Cre (bottom) mice injected with an INTRSECT Con/Fon-EYFP virus. Quantification (right) shows no EYFP expression, confirming dual recombinase dependence. **b**, Representative photomicrographs showing TH and mCherry immunoreactivity in TH-Flp (top) and VGLUT2-Cre (bottom) mice injected with an INTRSECT Coff/Fon-mCherry virus. Quantification (right) shows robust mCherry expression in TH-Flp mice but no expression in VGLUT2-Cre mice, consistent with Flp-dependent activation and Cre-dependent suppression. **c**, Confocal images (left) and 3D renderings (right) showing VGLUT2 (Slc17a6) transcript puncta (blue) within EYFP+ neurons (green; DA–GLU) and mCherry+ neurons (magenta; DA-only). **d**, Quantification of VGLUT2 transcript puncta per neuron in EYFP+ versus mCherry+ populations shows that VGLUT2 (a glutamatergic marker) is enriched in EYFP+ neurons, whereas mCherry+ neurons exhibit little to no VGLUT2 signal, with only a few cells containing a single punctum.

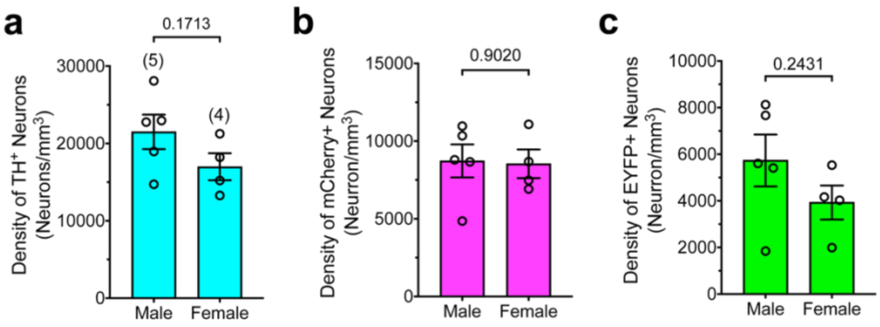


**Extended Fig. 2 | No sex differences in dopaminergic subpopulations density in the VTA of young mice.** Bar graphs comparing the density (neurons/mm³) of TH+ neurons (**a**), mCherry+ neurons (**b**), and EYFP+ neurons (**c**) in male versus female young mice. Each bar represents group mean ± SEM. P-values are displayed on the plots, indicating no statistically significant differences between sexes for any neuronal population shown.

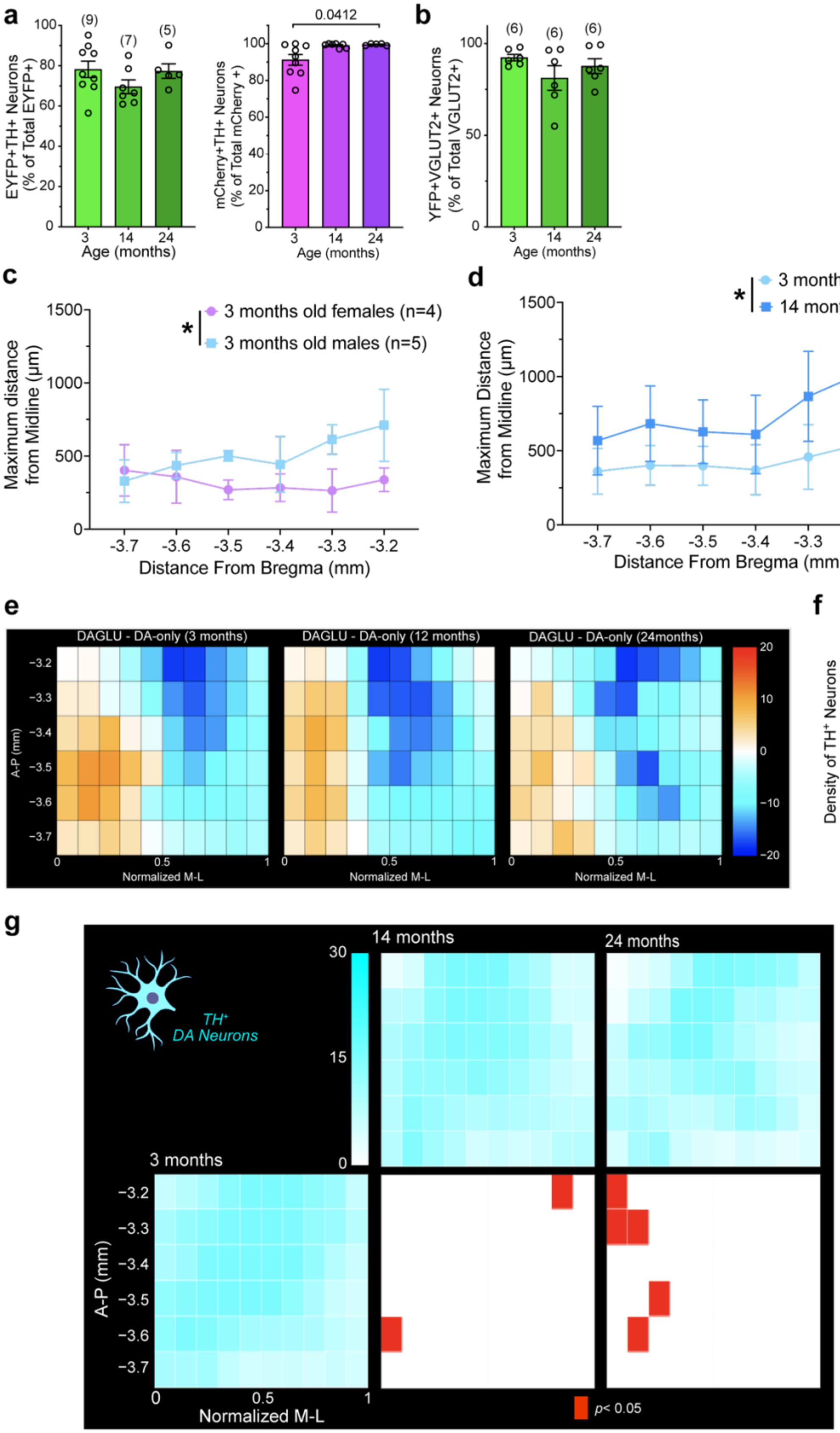

a
EYFP+TH+ Neurons (% of Total EYFP+)
(9) (7) (5)
Age (months)
mCherry+TH+ Neurons (% of Total mCherry +)
0.0412
b
YFP+VGLUT2+ Neurons (% of Total VGLUT2+)
(6) (6) (6)
c
3 months old females (n=4)
3 months old males (n=5)
Maximum distance from Midline (µm)
Distance From Bregma (mm)
d
3 months
14 months
Maximum Distance from Midline (µm)
e
DAGLU - DA-only (3 months)
DAGLU - DA-only (12 months)
DAGLU - DA-only (24months)
A-P (mm)
Normalized M-L
f
Density of TH+ Neurons (Neurons/mm3)
0.0093
0.0058
g
14 months
24 months
3 months
TH+ DA Neurons
A-P (mm)
Normalized M-L
p< 0.05

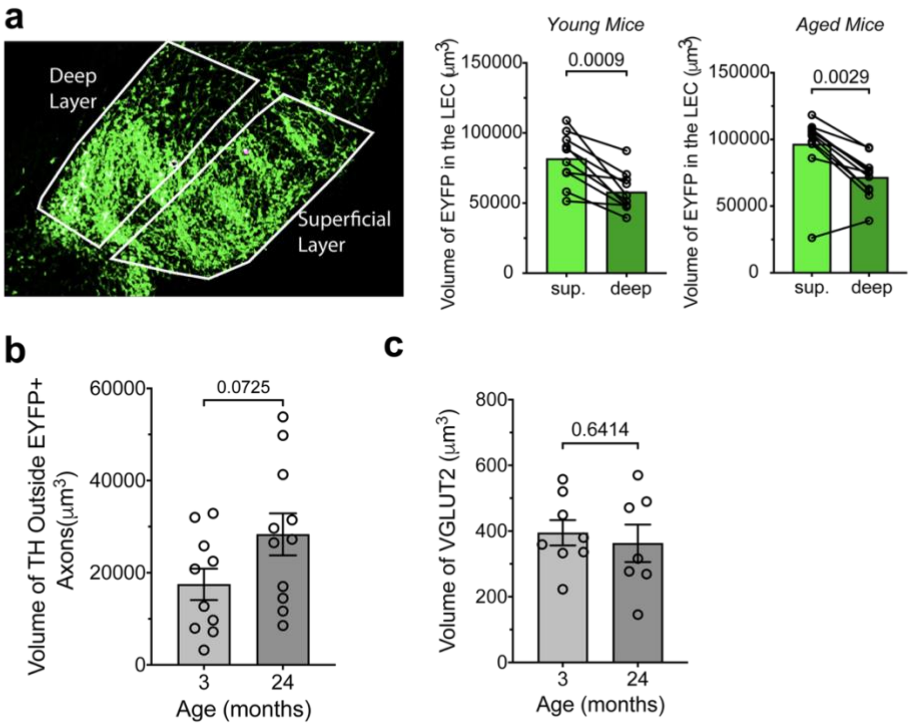


**Extended Fig. 4 | Layer- and age-dependent analysis of EYFP, TH, and VGLUT2 volume in the LEC. a**, Left, representative LEC image illustrating the delineation of superficial and deep layers for volumetric analysis of ChR2–EYFP labeled axons in DAT-IRES-Cre mice. Bar plots show total EYFP+ axonal volume (µm$^{3}$) in superficial versus deep layers at 3 (*left*) and 24 months (*right*). EYFP+ volume was greater in superficial than deep layers at both ages (3 months: paired t-test, $t(9)=4.882$, $P=0.0009$; 24 months: Wilcoxon matched-pairs signed-rank test, $W = -62$, $P=0.0029$). **b**, TH+ axonal volume (µm$^{3}$) outside EYFP+ axons in the LEC at 3 and 24 months; no age difference (unpaired t-test, $t(19)=1.876$, $P=0.0761$). **c**, Total VGLUT2+ terminal volume (µm$^{3}$) in the LEC at 3 and 24 months; no age difference (unpaired t-test, $t(13)=0.4768$, $P = 0.6414$). **d**, VGLUT2+ volume (µm$^{3}$) in superficial and deep layers at 3 and 24 months; no age effect (unpaired t-test, $t(13)=0.4768$, $P = 0.6414$). Each dot represents an animal; bars show mean ± SEM.

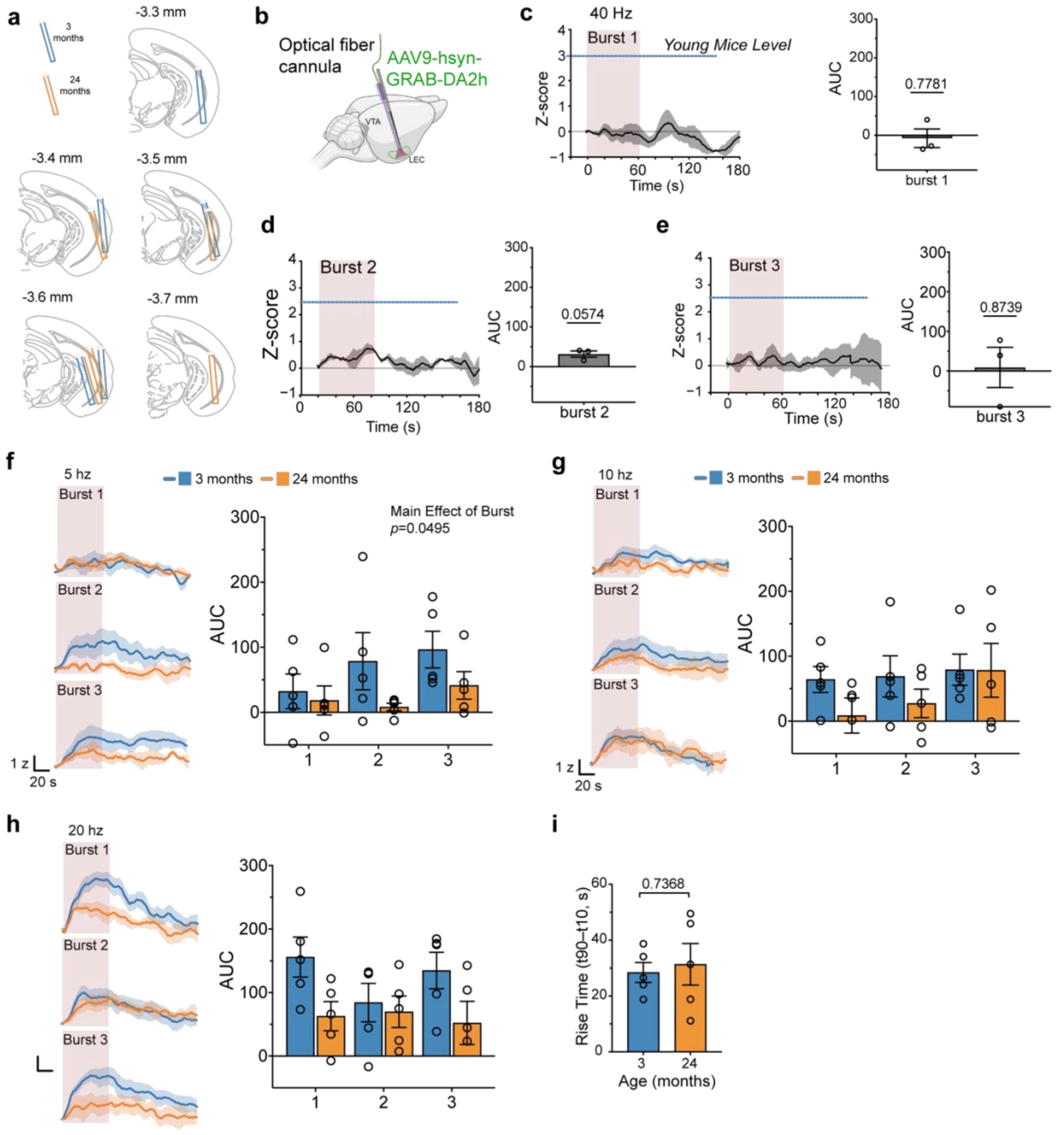


**Extended Fig. 5 | Validation of combined optogenetics and fiber photometry and dopamine responses to 5–20 Hz stimulation. a,** Schematic of optic fiber cannula placement in young and aged mice. **b**, Schematic of injection of $GRAB_{DA2h}$ virus and cannula placement for control experiments testing the effect of red-light stimulation on $GRAB_{DA}$ signals. **c–e**, Peri-event histograms showing mean $GRAB_{DA}$ responses to 40 Hz stimulation delivered as three consecutive 60-s bursts (n = 3 mice; 2 young, 1 aged; shaded area, ± SEM). Right, quantification of red-light effects on $GRAB_{DA}$ signals for bursts 1–3; one-sample t-tests showed no significant effect (burst 1: t(2)=0.3218, P=0.7781; burst 2: t(2)=3.991, P=0.0574; burst 3: t(2)=0.1797, P=0.8739). **f**, Mean traces (young, blue; aged, orange; shaded area, ± SEM) and quantification for 5 Hz stimulation across three consecutive bursts. Two-way repeated-measures ANOVA showed no main effect of age and no Age × Burst interaction, but a main effect of burst (Age × Burst: F(2,16)=1.607, P=0.2312; Burst: F(2,16)=3.648, P=0.0495; Age: F(1,8)=1.935, P=0.2016). **g**, Same as f for 10 Hz stimulation; no significant effects (Age × Burst: F(2,16)=0.6000, P=0.5607; Burst: F(2,16)=1.428, P=0.2826; Age: F(1,8)=1.432, P=0.2657). **h**, Same as f for 20 Hz stimulation; no significant effects (Age × Burst: F(2,16)=2.824, P=0.0890; Burst: F(2,16)=1.620, P=0.2287; Age: F(1,8)=3.222, P=0.1104). **i**, Rise time (time to 90% peak minus time to 10% peak) did not differ by age (unpaired t-test, t(8)=0
